# Mechanism of activation of an ancestral TEC kinase by PIP_3_

**DOI:** 10.1101/2025.05.22.653117

**Authors:** Eva Krötenheerdt, Lucas Piëch, Freia von Raußendorf, Ronja Reinhardt, Nicholas Wedige, Hunter G. Nyvall, Emma E. Walsh, John E. Burke, Thomas A. Leonard

## Abstract

The TEC kinases are a family of five paralogous mammalian genes that play crucial roles in cell growth, proliferation and differentiation, particularly in immune cells. The recruitment and activation of the TEC kinases depend on the generation of the lipid second messenger, PIP_3_, in the plasma membrane. However, the mechanisms by which PIP_3_ activates the TEC kinases are not well understood. We have elucidated the autoinhibited conformation of an ancestral TEC kinase from the choanoflagellate *Monosiga brevicollis.* We demonstrate that PIP_3_ relieves autoinhibition of MbTEC by displacing its PH domain from an evolutionarily conserved inhibitory interaction with its kinase domain. We also show that a conserved polyproline motif within MbTEC promotes its activation in a kinase-intrinsic mechanism. Finally, we show that the PH domain is sufficient to restore autoinhibition in a constitutively active mutant of MbTEC. Our findings reveal that PIP_3_ is necessary and sufficient for both MbTEC activation and inactivation.

**Significance Statement:** The Tec family of protein kinases plays an essential role in cell signaling, particularly in the proliferation and differentiation of immune cells. Consequently, their dysregulation is causative of inherited immunodeficiency, while the Tec kinases are also therapeutic targets in the control of hematological malignancies. We have elucidated a conserved mechanism by which the Tec kinases are activated by the lipid second messenger PIP_3_. PIP_3_ is necessary and sufficient for Tec activation, while its turnover is sufficient for Tec inactivation. Our work identifies PIP3 as the ultimate gatekeeper of Tec activity in cells, with implications for the rationalization and treatment of human disease.

## Introduction

The TEC kinases are a subfamily of the Src family of non-receptor tyrosine kinases (SFK), best known for their roles in the proliferation and differentiation of immune cells (1). Mutations in Bruton’s tyrosine kinase (BTK), one of the five mammalian TEC kinase paralogs, are associated with X-linked agammaglobulinemia (XLA), a disease characterized by a failure of pre-B cells to mature into terminally differentiated, antibody-producing B cells and a consequent lack of circulating antibodies (2). Mutations in the same gene lead to X-linked immunodeficiency (XID) in mice (3). Conversely, inhibition of BTK is used in the clinic to treat a range of B cell malignancies, including chronic lymphocytic leukemia (CLL), mantle cell lymphoma, and Waldenström’s macroglobulinemia (4).

The TEC kinases are distinguished by an N-terminal pleckstrin homology (PH) domain, which is appended to an ancient Src homology 3 (SH3)-Src homology 2 (SH2)-kinase (32K) module shared by all SFKs (5). BTK is autoregulated by its SH3 and SH2 domains, which pack against the kinase domain to maintain it in an inactive conformation (6, 7), as also observed for SRC, hematopoietic cell kinase (HCK) and Abelson murine leukemia viral oncogene homolog 1 (ABL) (8–10). The SH2 and SH3 domains bind to phosphorylated tyrosine marks and polyproline motifs in activated receptors at the plasma membrane, respectively. This leads to the specific recruitment of BTK to membrane proximal sites, which is a prerequisite for its activation and subsequent downstream substrate phosphorylation. Early reports suggested that the activation loops of BTK and inducible T-cell kinase (ITK) are *trans*-phosphorylated by other SFKs (11–13), while more recent reports have attributed activation loop phosphorylation to autocatalysis (6, 7, 14).

Unlike other SFKs, which are localized to the plasma membrane via myristoylation of their N-termini (15), the localization of the TEC kinases depends on the lipid second messenger phosphatidylinositol-3,4,5-trisphosphate (PIP_3_). PIP_3_ produced by class I phosphatidylinositol 3-kinases (PI3K) immediately downstream of the B cell receptor (BCR) or T cell receptor (TCR) is required for re-localization of BTK and ITK, respectively, from the cytosol to the plasma membrane. Various functions have been ascribed to the PH domains of BTK and ITK, from targeting to the plasma membrane (16–18) to PIP_3_-mediated dimerization (6, 14, 19) and autoinhibition of kinase activity (6, 7, 20–22). While membrane targeting and co-localization of ITK with the TCR can be substituted with the membrane-targeting myristoylation sequence of LCK, the PH domain is indispensable for kinase activation (23). Although full-length BTK has recently been crystallized, the PH domain was not resolved in the electron density map (7), while studies of BTK in solution and by cryo-electron microscopy have concluded that the protein adopts a spectrum of conformations with varying degrees of compaction (7, 24, 25). Together with the available experimental structures, mapping of the conformation of the PH domain in BTK and ITK by hydrogen-deuterium exchange mass spectrometry (HDX-MS) and nuclear magnetic resonance (NMR) spectroscopy has led to proposals of an ensemble of conformational states in solution (7, 20, 21, 26). The role, therefore, of the PH domain in regulating the activity of the TEC kinases is still incompletely understood.

The TEC kinases first emerged in unicellular choanoflagellates, before gene duplication, prior to the evolution of vertebrates, gave rise to the five mammalian paralogs (27). Tissue-specific expression patterns, particularly in immune cells, have since given rise to the diversification of the signaling pathways regulated by individual paralogs (28). To elucidate the mechanism by which the TEC kinases are regulated by PIP_3_, we studied an ancestral TEC kinase from the choanoflagellate *Monosiga brevicollis*, which we hereinafter refer to as MbTEC. We show that MbTEC adopts a compact conformation in solution and that the PH domain both stabilizes this conformation and inhibits its kinase activity. We provide evidence that the PIP_3_-binding pocket of the PH domain is sequestered in a specific autoinhibitory conformation centered on the kinase domain of MbTEC and that mutation of either side of the PH-kinase interface activates MbTEC independently of PIP_3_. Finally, we demonstrate that a conserved polyproline motif in MbTEC plays a crucial role in its allosteric, PIP_3_-mediated activation. Our findings reveal that PIP_3_ is both necessary and sufficient for MbTEC activation and inactivation, thereby confining TEC activity to the membrane. TEC kinases are, therefore, coincident detectors of PIP_3_, phospho-tyrosine and polyproline motifs, all of which must be engaged for the recruitment and activation of TEC with high fidelity.

## Results

### *Monosiga brevicollis* TEC adopts a conserved autoinhibited conformation

MbTEC is a 667 amino acid long protein with 42% sequence identity to human BTK. It comprises a PH domain at its N-terminus followed immediately by a TEC homology (TH) domain, an unstructured linker of approximately 85 amino acids that contains a conserved polyproline motif, SH3, SH2 and kinase domains (Figure 1A). The TH domain is not an independently folding domain, but a zinc finger that is an intrinsic part of the PH domain fold (16). MbTEC Y559 in the activation loop corresponds to the canonical autophosphorylation site (human BTK Y551). A conserved hydrophobic amino acid in the SH2-kinase linker (MbTEC L396) forms the central component of a three-residue hydrophobic stack (Figure 1B-D) that serves to maintain the SH3-SH2-kinase module of SFKs in an inactive conformation stabilized by ADP (29). Mutation of the central residue in the hydrophobic stack is sufficient to convert both SRC and ITK into an active conformation, concomitant with the binding of ATP (29).

**Figure 1.**
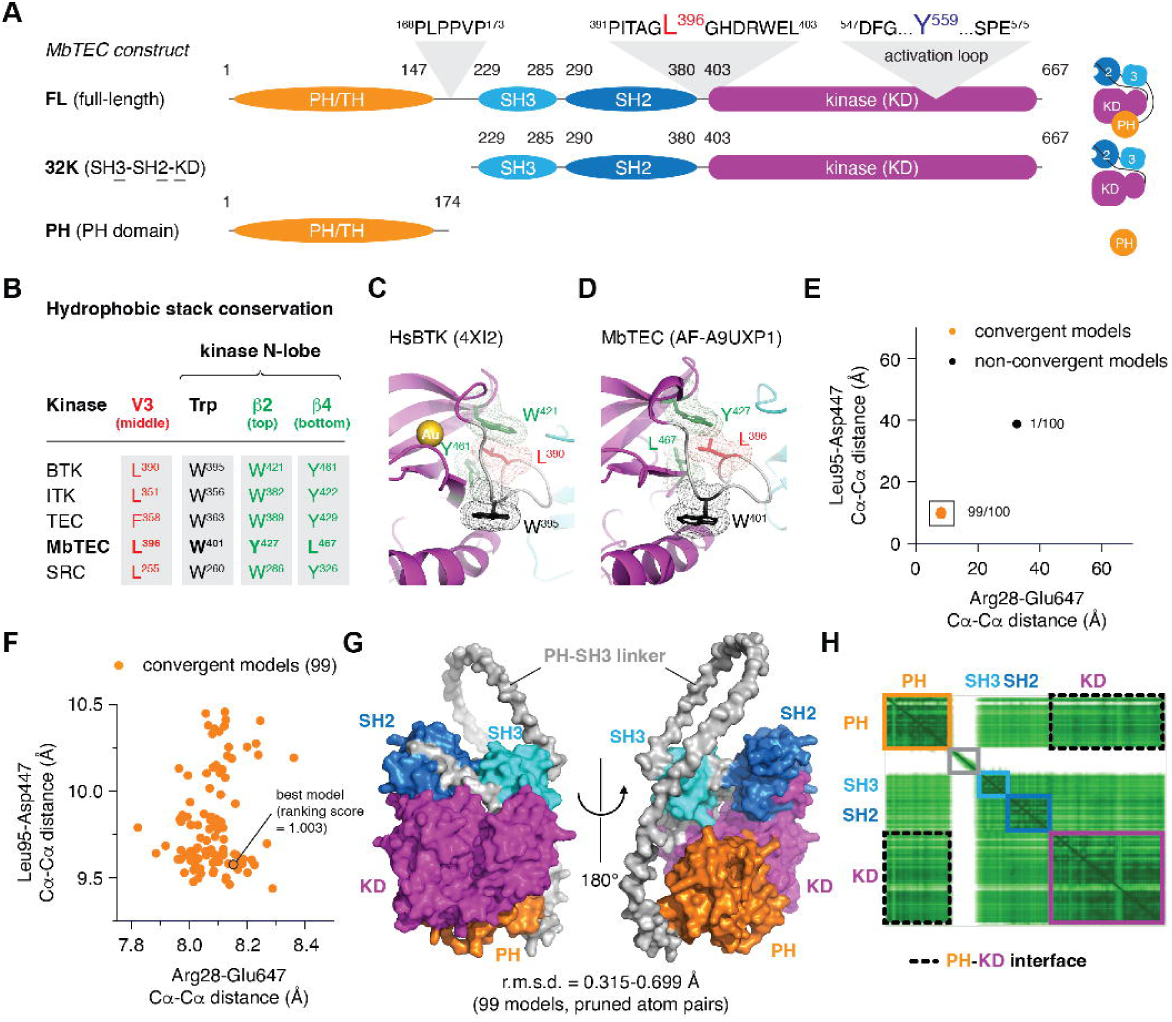
*M. brevicollis* TEC adopts a conserved autoinhibited conformation. A. Domain schematics of all protein constructs used in this study. B. Hydrophobic stack conservation in the TEC kinases. C. Structure of hydrophobic stack of mouse BTK. The middle residue in the stack, contributed by the SH2-kinase linker, is shown in red. The three residues in the stack are shown in mesh representation. D. AlphaFold-predicted structure of MbTEC hydrophobic stack. The three residues in the stack are shown in mesh representation. E. Euclidean distance plot of two PH-kinase domain residue pairs in 100 models generated by AlphaFold 3 from a curated, taxonomically balanced MSA. Inter-residue Cα-Cα distances for Arg28-Glu647 and Leu95-Asp447 provide an indicator of model convergence. F. Zoomed in plot of the 99 convergent models shown in the black box in panel E. The best ranked model is indicated with a black outline and its AlphaFold 3 ranking score. G. Overall structure of the best ranked model and root mean square deviation over pruned atom pairs between the 99 models. H. Pair alignment error plot for the best ranked model. Individual domains are depicted with colored rectangles according to the color scheme indicated in panel A. Black, dashed rectangles indicate the regions of the plot that correspond to residue pairs between the PH and kinase domains.

To study the properties of MbTEC, we purified wild-type and mutant variants of the full-length protein as well as its isolated PH domain and 32K modules (Figure 1A) to homogeneity. All kinase domain-containing constructs were dephosphorylated *in vitro* with the tyrosine phosphatase YopH and the serine/threonine lambda phosphatase after affinity purification, prior to ion exchange and size-exclusion chromatography. The mass and polydispersity of all purified proteins employed in this study were confirmed by mass spectrometry (Supplementary Figure 1A-N) and mass photometry (Supplementary Figure 2A-L) or size-exclusion chromatography coupled to multi-angle light scattering (SEC-MALS) (Supplementary Figure 2C). To investigate the nucleotide dependence of MbTEC, we measured its thermal stability in the presence of ADP, ATP, and a non-hydrolyzable ATP analog, AMPPNP. MbTEC was preferentially stabilized by ADP (Supplementary Figure 3A), consistent with what has previously been observed for SRC and ITK (29).

Advances in computational modeling have made the determination of proteins of unknown structure routine (30), including, recently, the docking of ligands and ions in AlphaFold 3 (31). Using a standard implementation of AlphaFold 3, we observed consistent model convergence on the known intramolecular architecture of the SH3, SH2 and kinase domains, but divergent placement of the PH domain (Supplementary Figure 3B). Suspecting that AlphaFold may be failing to detect a co-evolutionary signal for the PH-kinase interface in the multiple sequence alignment (MSA) used for structure inference, we investigated the quality of the MSA. AlphaFold 3 uses JackHMMER (32) to detect remote sequence homologs of protein domains using profile hidden Markov models such that evolutionary sequence depth provides strong couplings that guide structure inference (32). However, an MSA that contains partially homologous sequences that do not belong to the same family will dilute physiologically relevant evolutionary couplings between domains. For example, tyrosine kinases that belong to the Src superfamily, but do not possess a PH domain, will not exhibit evolutionary couplings corresponding to an intramolecular interaction with the PH domain. Equally well, taxonomic bias, such as overrepresentation of chordate sequences, will dominate the MSA and potentially mask evolutionary covariance, affecting the structure inference step.

To address these problems, we manually curated a set of 2,442 TEC kinase sequences by filtering UniProt for the presence of PH, Btk_Zn_finger, SH3, SH2, and tyrosine kinase domains as defined by InterPro (33) (Supplementary Figure 3C). After aligning the sequences using MAFFT (34, 35), we corrected for taxonomic bias by reweighting or downsampling (Supplementary Methods) and fed the resulting MSAs into AlphaFold 3. To determine model convergence, we selected two pairs of residues between each of the regulatory PH, SH3 and SH2 domains and the kinase domain and plotted the Euclidean distances between the Cα atoms of each pair of residues (Supplementary Figure 3D). In the standard AlphaFold 3 pipeline without MSA curation, 39% model convergence with an identical PH-kinase domain interface was obtained when including a single Zn^2+^ ion (Supplementary Figure 3D-E), which is present in an experimentally determined structure of the PH domain in BTK (16). Structure prediction with our curated MSA resulted in a modest increase in model convergence to 52% (Supplementary Figure 3C), while taxonomic reweighting of the sequences increased this figure to 80% (Supplementary Figure 3C) and taxonomic downsampling increased this further to 99% (Figure 1E-F, Supplementary Figure 3C, F-H). The resulting MbTEC structures exhibit a conserved 32K architecture superimposable with the crystal structure of the 32K module of human Btk (6) and the ADP-bound conformation of Src (29), as well as structures of Hck (8) and Abl (10) (Supplementary Figure 3I-J). All models capture the inactive conformation of the activation loop tyrosine, which is sequestered in a non-phosphorylatable position. Finally, the pair-alignment error (PAE) plot for the highest scoring (AlphaFold 3 ranking score) model (Figure 1F-G) exhibits strong residue couplings between the PH and kinase domains (Figure 1H). These findings imply an evolutionarily conserved interaction between PH and kinase domains in the TEC kinases.

### MbTEC adopts a compact conformation in solution

To characterize the size, shape and conformation of MbTEC in solution, we employed size exclusion chromatography coupled to small-angle X-ray scattering (SEC-SAXS). Frames were averaged across the elution peak and reduced to a 1D scattering curve (Figure 2A). Guinier plots for MbTEC^FL^ and MbTEC^32K^ are both linear in the low-angle regime (Supplementary Figure 4A-B), indicating that the radius of gyration is well estimated in each case. From the pair distribution functions (Figure 2B), MbTEC^FL^ has an *R*_g_ of 3.1 nm and a maximum dimension of 8.4 nm compared with 2.5 nm and 8.3 nm for MbTEC^32K^ (Figure 2C). Estimation of the molecular mass from the scattering is within 2 kDa of the actual mass of the proteins (Figure 2C). Calculation of the theoretical solution scattering of our AlphaFold 3 model for the autoinhibited conformation of MbTEC indicated a remarkable fit to the experimental scattering, both for MbTEC^FL^ and MbTEC^32K^ (Figure 2A). *Ab initio* calculation of the respective molecular envelopes from the scattering curves and fitting of the respective models for MbTEC^FL^ (Figure 2D) and MbTEC^32K^ (Figure 2E) also revealed a strong correlation between model and experiment. This suggests MbTEC forms a compact conformation in solution that is well described by the AlphaFold model in Figure 1G.

**Figure 2.**
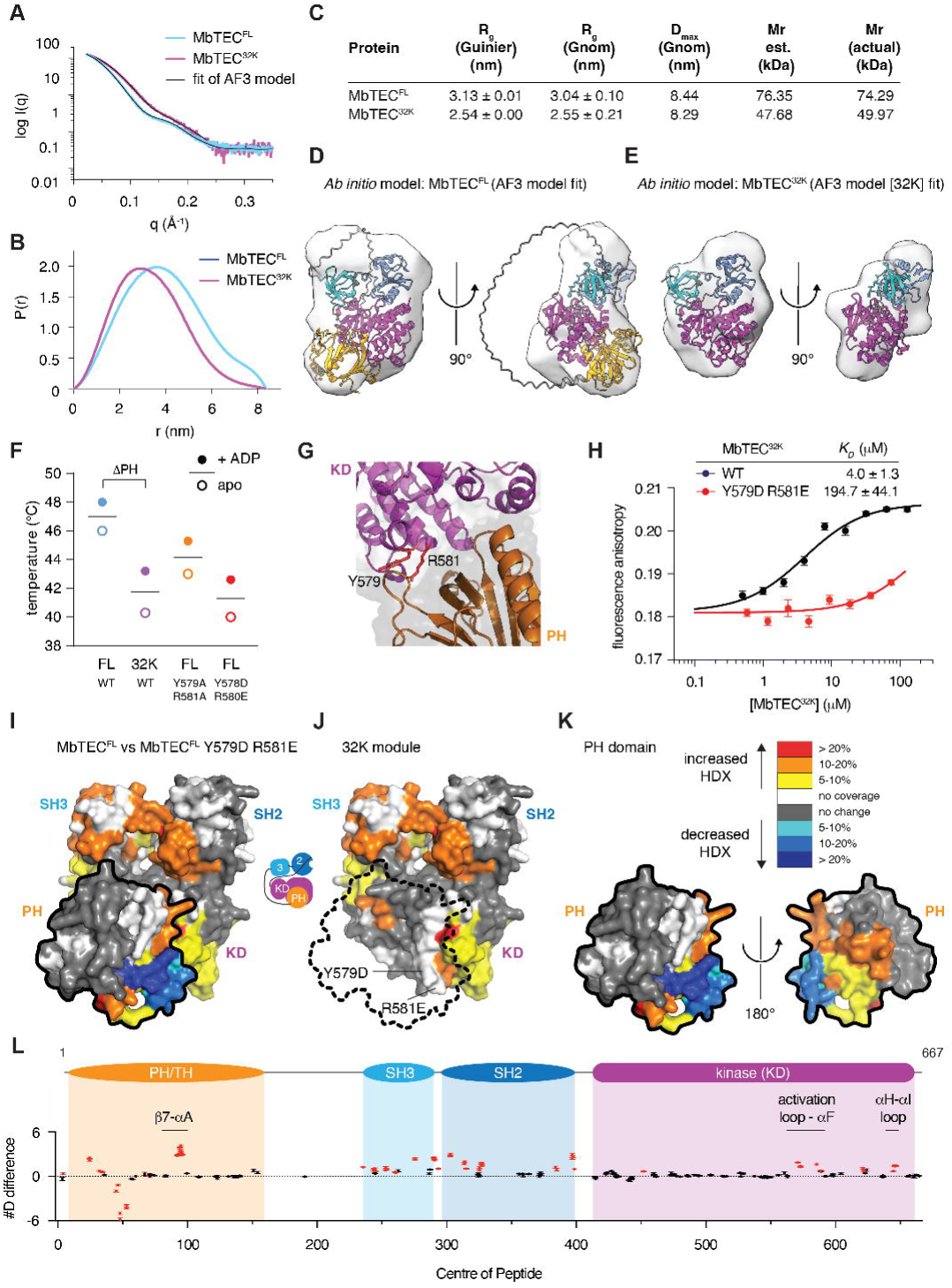
MbTEC adopts a compact conformation in solution. A. Small-angle X-ray scattering (SAXS) curves of MbTEC^FL^ and MbTEC^32K^ in solution. The theoretical scattering curves of the AlphaFold models of MbTEC^FL^ and MbTEC^32K^ and the fits to their respective scattering curves are shown as solid lines. B. Pair distribution functions for MbTEC^FL^ and MbTEC^32K^. C. Summary table of particle parameters derived from SAXS measurements. D. *Ab initio* molecular modeling of the molecular envelope of MbTEC^FL^. The AlphaFold model of MbTEC^FL^ was docked into the envelope using SUPCOMB (57). E. *Ab initio* molecular modeling of the molecular envelope of MbTEC^32K^. The AlphaFold model of MbTEC^32K^ was superimposed with the model using SUPCOMB (57). F. Melting temperatures of MbTEC^FL^ (blue), MbTEC^32K^(purple), MbTEC^FL^ Y579A R581A(orange) and MbTEC^FL^ Y579D R581E (red) with (filled circle) or without (empty circle) ADP. G. Location of Y579 and R581 in the PH-kinase domain interface. H. Binding of MbTEC^32K^ (wt) and MbTEC^32K^ Y579D R581E to MbTEC^PH^ determined by fluorescence anisotropy. Each data point is the average of 10 measurements. The data were fit with a one-site binding model to determine *K*_D_. I. Pairwise HDX-MS analysis of MbTEC^FL^ compared to MbTEC^FL^ Y579D R581E. Significant differences in HDX in MbTEC^FL^ Y579D R581E are mapped onto the AlphaFold3 model of MbTEC^FL^ and are color coded according to the legend. The PH domain is outlined with a black stroke. J. HDX changes mapped on the 32K module of MbTEC (PH domain cut away). The surface of the PH domain is outlined with a dashed stroke. The location of Y579 and R581 are indicated on the kinase domain. K. HDX changes mapped on the PH domain of MbTEC. L. Sum of #D difference in deuterium incorporation over the entire time course. Each point represents and individual peptide with significant changes shown in red (greater than 0.40 Da, 5% difference and two tailed t-test p<0.01 at any timepoint, n=3). Domain architecture is annotated above.

Consistent with the loss of an energetically favorable interface, deletion of the PH domain resulted in a 6°C reduction in thermal stability (Figure 2F, Supplementary Figure 4C). Mutation of Y579 and R581, conserved residues in the interface (Figure 2G) to alanine reduced thermal stability by 3°C, while their mutation to asparate and glutamate respectively resulted in the same thermal stability as MbTEC^32K^ lacking its PH domain (Figure 2F, Supplementary Figure 4D). These observations indicate that substitution of Y579 and R581 with alanine weakens the autoinhibitory conformation by reducing van der Waals contacts, but substitution with charged residues that introduce unfavorable interactions is sufficient to completely disrupt the interface. Consistently, MbTEC^32K^ bound to the PH domain with an affinity of 4.0 μM, but binding of MbTEC^32K^ Y579D R581E was barely detected (Figure 2H).

To map the intramolecular association of the MbTEC PH and kinase domains experimentally, we employed hydrogen-deuterium exchange mass spectrometry (HDX-MS). HDX-MS provides peptide-level information on the rate of exchange of backbone amide hydrogens with a deuterated solvent, thereby revealing changes in secondary structure when comparing two species. To investigate the surfaces of MbTEC involved in PH domain sequestration, we compared the rates of HDX between MbTEC^FL^ and MbTEC^FL^ Y579D R581E. Mutation of the interface resulted in significant increases in the rate of HDX throughout the SH3 domain, SH2 domain, PH domain and in the αC helix, activation loop, and αH-αI loop of the kinase domain (Figure 2I-L). There was a single region in the PH domain that showed decreased exchange in the mutant. These changes could be recapitulated by deletion of the PH domain (Supplementary Figure 4E-H), HDX increases similar to those previously reported for the same comparison in human BTK (20). The αC helix, activation loop, and αH-αI loop are all predicted to be at least partially sequestered in interactions with the PH domain. This data also suggests a role of the PH domain in stabilizing inhibitory contacts of the SH2 and SH3 domain. The time course of deuterium incorporation is shown for individual peptides in these regions in Supplementary Figure 4I (MbTEC^FL^ vs MbTEC^32K^) and Supplementary Figure 4J (MbTEC^FL^ vs MbTEC^PH^), with the full set of deuterium incorporation data shown in the source data.

### MbTEC is autoinhibited by its PH domain

Although the activation of TEC kinases by PIP_3_-containing membranes has previously been demonstrated (6, 7, 14, 21, 26), these experiments cannot deconvolute the relative influences of local concentration and allosteric effects on TEC activation. Since autophosphorylation depends on dimerization, the local concentration of a TEC kinase on a membrane will necessarily result in its activation, regardless of whether there is an allosteric component. Other studies, however, have reported an increase in BTK and ITK activity upon deletion of the PH domain (6, 22) as well as conformational changes in BTK upon PIP_3_ binding (26), indicating that the PH domain exerts an autoinhibitory effect in the absence of PIP_3_. To determine whether the PH domain exerts an autoinhibitory effect on MbTEC, we compared the rates of autophosphorylation of MbTEC^FL^, MbTEC^FL^ L396A, MbTEC^32K^ and MbTEC^32K^ L396A using a radiometric kinase assay (Figure 3A). MbTEC^FL^ exhibited the lowest rate of autophosphorylation, which was only modestly increased by either deletion of the PH domain or mutation of L396 in the SH2-kinase linker, a mutation known to promote the constitutive activation of ITK (29). Combining deletion of the PH domain with mutation of L396, however, increased the rate of autophosphorylation more than 10-fold (Figure 3A). These observations indicate that the PH domain exerts a strong autoinhibitory influence on kinase activity that cannot be overcome by disruption of the SH3-kinase interaction.

**Figure 3.**
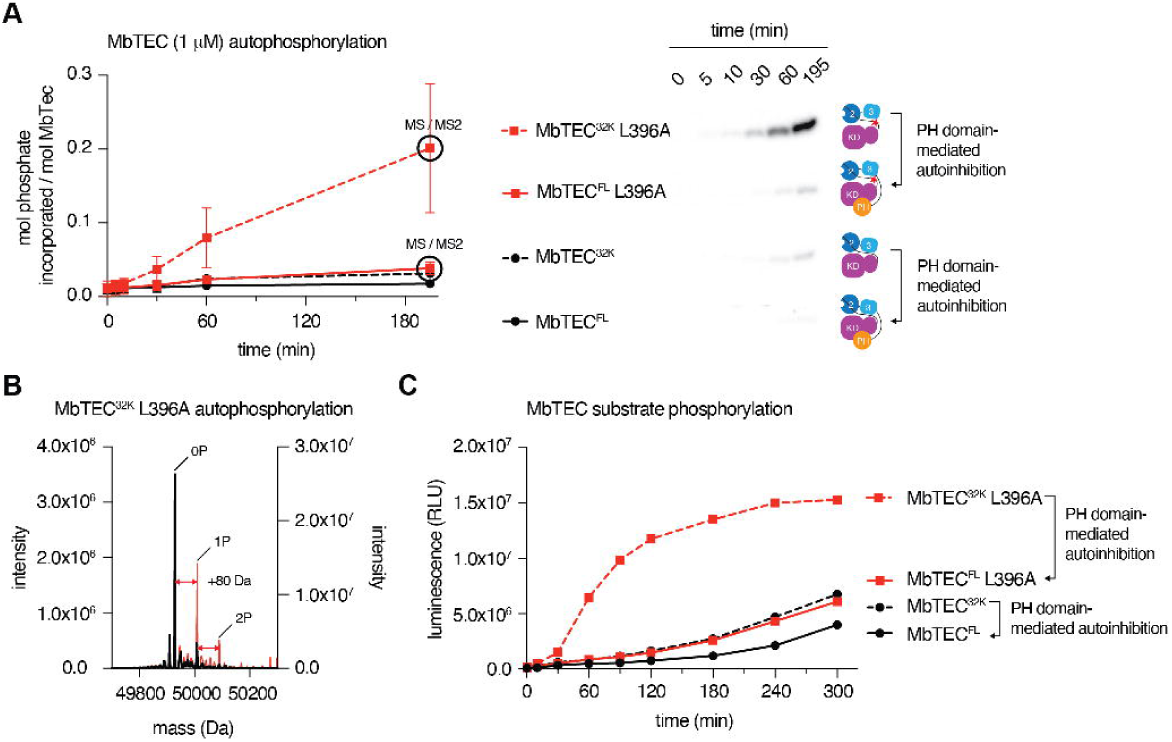
MbTEC is autoinhibited by its PH domain. A. Radiometric autophosphorylation assays of MbTEC^FL^ (black, solid line) and MbTEC^32K^ (black, dashed line), as well as their L396A counterparts (red). Right: representative cartoons and autoradiograph for each construct. B. Mass spectrometry analysis of the product of MbTEC^32K^ L396A autophosphorylation (denoted by the black circle in panel A). Intensities before autophosphorylation in black (left axis) and after autophosphorylation in red (right axis). C. Substrate peptide phosphorylation by MbTEC^FL^ (black, solid line), MbTEC^FL^ L396A (black, dashed line), MbTEC^32K^ (red, solid line) and MbTEC^32K^ L396A (red, dashed line). ADP-Glo kinase assay; RLU (raw luminescence units).

To assess the stoichiometry of autophosphorylation of MbTEC, we measured the mass of MbTEC^FL^ L396A and MbTEC^32K^ L396A pre- and post-autophosphorylation by mass spectrometry. We analyzed samples in the linear kinetics regime to capture the most specific events. The increase in mass corresponded to modification with a single phosphate in 40% of MbTEC^32K^ L396A molecules under the conditions of the assay (Figure 3B). By contrast, MbTEC^FL^ L396A exhibited negligible autophosphorylation (Supplementary Figure 5A), consistent with robust autoinhibition by its PH domain. To identify the modified residue, MbTEC^32K^ L396A and MbTEC^FL^ L396A were subjected to tandem mass spectrometry, which revealed phosphorylation of the canonical activation loop tyrosine Y559 in MbTEC^32K^ L396A, but not MbTEC^FL^ L396A (Supplementary Figure 5B). This confirmed that the PH domain specifically inhibits activation loop phosphorylation of MbTEC.

Finally, we observed that phosphorylation of a consensus substrate peptide is also robustly activated by the additive effects of both PH and SH3 domain displacement (Figure 3C). Together, these observations indicate that MbTEC is robustly autoinhibited by its PH domain, and that PH domain displacement is necessary for its activation.

### A gain-of-function mutation in the PH domain activates MbTEC

A previously identified gain-of-function mutation in the PH domain of BTK, E41K, has been proposed to activate BTK by enhancing its binding to PIP_3_ at the plasma membrane (17, 18, 36) as well as promoting IP_6_-mediated dimerization (37). Interestingly, E41 is predicted to make hydrogen bonds to R28 and K53 in the autoinhibited conformation of MbTEC (Figure 4A), where R28 is also stabilized by a hydrogen bond to E647 in the kinase domain. As such, the E41K mutation would be expected to destabilize the PH-kinase interface by introducing charge repulsion with R28 and K53. Consistent with this hypothesis, MbTEC^FL^ E41K (equivalent to human BTK E41K) exhibited a significantly increased rate of autophosphorylation in the absence of PIP_3_ compared to wild-type MbTEC^FL^, equivalent to the rate of autophosphorylation of MbTEC^32K^ L396A, indicating that the E41K mutation weakens PH domain-mediated autoinhibition (Figure 4B). The kinetics of MbTEC^FL^ E41K autophosphorylation could be further enhanced by introduction of the L396A mutation, simulating SH3 domain release (Figure 4B). Although R29X in human BTK is a loss-of-function mutation in XLA due to a loss of PIP_3_ binding and a consequent failure to activate BTK downstream of the BCR (16, 17, 38), mutation of MbTEC^PH^ R28 to glutamate also weakened the PH-kinase interface by 5-fold (Figure 4C), consistent with the sequestration of the PIP_3_ binding pocket predicted by our model.

**Figure 4.**
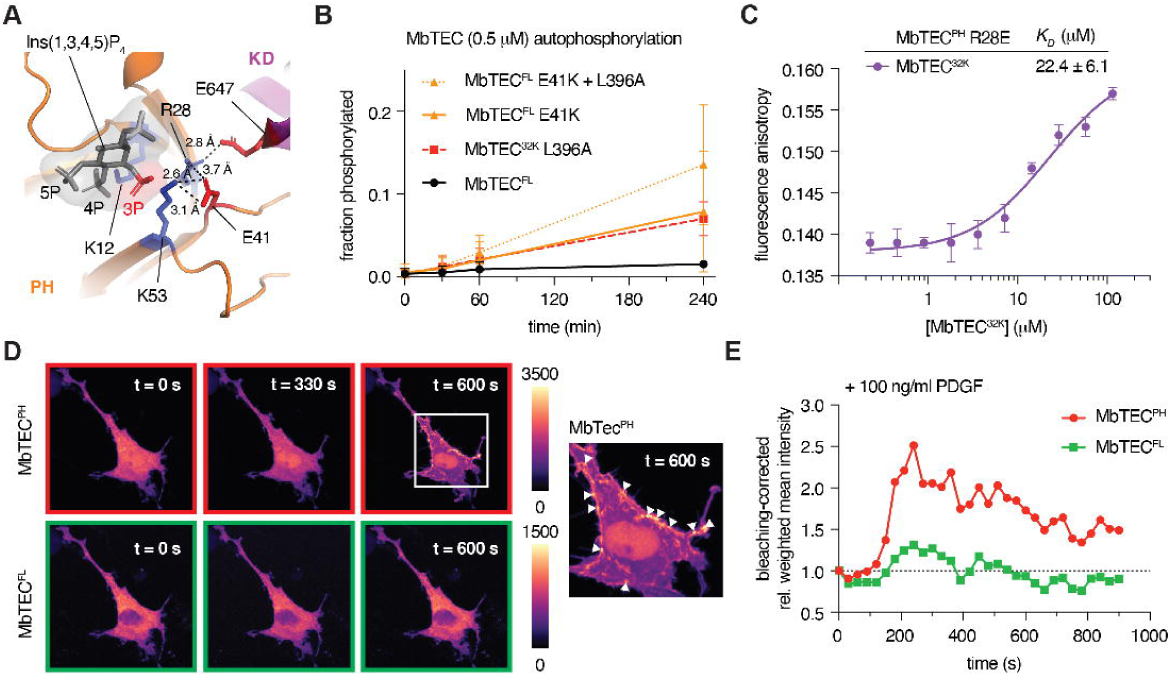
A gain-of-function mutation in the PH domain activates MbTEC. A. High-resolution view of the PIP_3_ binding site in the context of autoinhibited MbTEC. Residues that contribute to both PIP_3_ recognition and stabilization of the PH-kinase interface are depicted in sticks colored according to their acidic (red) or basic (blue) character. The head group of PIP_3_ was positioned in the model by superimposing the structure of the BTK PH domain in complex with Ins(1,3,4,5)P_4_ (PDB: 1B55). The 3’ phosphate crucial for TEC activation is colored red. Hydrogen bonds are indicated with dashed lines. B. Radiometric autophosphorylation assays of MbTEC^FL^ (black, solid line) and MbTEC^FL^ E41K (orange, solid line), as well as MbTEC^32K^ L396A (red, dashed line) and MbTEC^FL^ E41K L396A (orange, dashed line). Measurements are the average of three independent replicates. C. Binding of MbTEC^32K^ (wt) to MbTEC^PH^ (R28E) determined by fluorescence anisotropy. Each data point is the average of 10 measurements. The data were fit with a one-site binding model to determine *K_D_*. D. Representative images of the membrane translocation of MbTEC^FL^ (green channel) versus MbTEC^PH^ (red channel) over a time course of 10 min in co-transfected, PDGFαβ-treated NIH3T3 fibroblasts. Right: lookup tables (LUT) representing the color-encoded intensity map of individual pixels. White arrowheads: membrane foci of enriched MbTEC^PH^. E. Bleaching-corrected, normalized mask intensity profiles of MbTEC^FL^ (green) and MbTEC^PH^ (red) in a representative cell as a function of time, post-PDGFαβ stimulation.

Partial sequestration of the PIP_3_-binding surface by the kinase domain implies that there is a competition between PIP_3_ and the kinase domain for binding to the PH domain. The logical consequence of this should be a reduced affinity for PIP_3_ in full-length MbTEC and, conversely, enhanced binding by the isolated PH domain. To test this hypothesis in cells, we compared the membrane translocation behavior of co-expressed MbTEC^FL^ with MbTEC^PH^ within individual growth factor stimulated fibroblasts. MbTEC^FL^ exhibited a primarily diffuse cytoplasmic distribution with minor enrichment of MbTEC^PH^ at membrane protrusions (Figure 4D). Upon stimulation of the cells with PDGFαβ, MbTEC^PH^ exhibited considerably stronger enrichment at membrane protrusions in comparison to MbTEC^FL^ (Figure 4D-E, Supplementary Figure 6A). Importantly, while the membrane association of MbTEC^FL^ was transient, MbTEC^PH^ exhibited prolonged association with the membrane long after MbTEC^FL^ had returned to its basal level post-stimulation of the cells with PDGFαβ (Figure 4E, Supplementary Figure 6A). Together, these observations support an autoinhibitory role for the PH domain in which the PIP_3_-binding pocket is partially sequestered in the interface and PIP_3_ elicits MbTEC activation by destabilizing a network of electrostatic interactions between the PH domain and the kinase domain.

### MbTEC is allosterically activated by PIP_3_

Structurally and biochemically, the evidence presented so far strongly supports the allosteric activation of MbTEC by PIP_3_. To demonstrate it directly, we employed PIP_3_-containing lipid nanodiscs (Figure 5A). We observed that the rate of MbTEC autophosphorylation is dramatically enhanced by PIP_3_ binding (Figure 5B-C). Mass spectrometry revealed autophosphorylation of up to eight sites in MbTEC elicited by PIP_3_ (Supplementary Figure 7A), of which five sites correspond to tyrosines in the SH3 and SH2 domains (Supplementary Figure 7B-C), as well as *trans*-phosphorylation of the nanodisc membrane scaffold protein (MSP1D1) (Supplementary Figure 7D). These observations indicate that additional interactions within the ‘signalosome’ are likely required for high-fidelity auto- and substrate phosphorylation.

**Figure 5.**
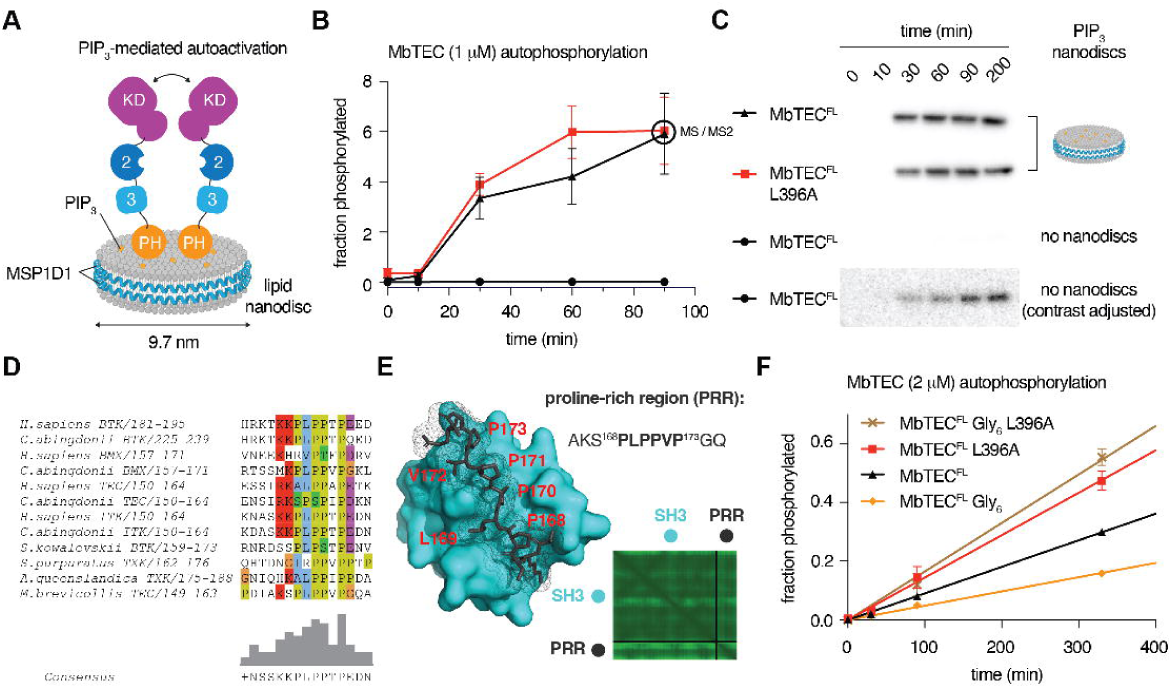
MbTEC is allosterically activated by PIP_3_. A. PIP_3_-doped nanodiscs used to activate MbTEC. MSP = membrane scaffold protein). B. PIP_3_-mediated activation of MbTEC^FL^ (black, triangles) and MbTEC^FL^ L396A (red, squares). Black, circles: MbTEC^FL^ without PIP_3_ nanodiscs. C. Representative autoradiographs for each construct. Note: the contrast has been artificially enhanced for the purposes of displaying the low basal autophosphorylation of MbTEC^FL^ in the absence of PIP_3_ nanodiscs. 200 min time points were not quantified in panel B due to saturation of the phosphor screen. D. Sequence conservation of the polyproline motif between TEC paralogs and over the course of vertebrate evolution. Representative human and turtle (*C. abingdonii*) sequences. E. AlphaFold prediction of the intramolecular association of the TEC kinase polyproline motif with its SH3 domain. F. Radiometric autophosphorylation time course of MbTEC^FL^ (black) and MbTEC^FL^ L396A (red), as well as MbTEC^FL^ Gly_6_ (orange) and MbTEC^FL^ Gly_6_ L396A (brown).

Curiously, an additive effect of PIP_3_ binding and displacement of the SH3 domain (L396A) was not manifested in increased kinetics of autophosphorylation (Figure 5B-C), as previously observed when the PH domain was deleted (Figure 3A). This finding suggests that PIP_3_ binding elicits complete allosteric activation of MbTEC^FL^, which cannot be phenocopied by simple deletion of the PH domain in MbTEC^32K^. Previous studies have reported an intramolecular association of the SH3 domain with a conserved proline-rich region (PRR) in the PH-SH3 interdomain linker (39–41) (Figure 1A, 5D), while a mutagenesis study has implicated the PRR in the activation of BTK (26). As proposed for BTK, the intramolecular association of the SH3 domain with this motif upon the allosteric activation of MbTEC by PIP_3_ is supported by AlphaFold predictions of the interface which converge on near-identical, high confidence models (Figure 5E).

These structures are superimposable with experimentally determined structures of the SH3 domain associated with the polyproline motif of both ITK (39) and TEC (40). To test whether the polyproline motif contributes to MbTEC activation in the absence of adaptor proteins, we compared the autophosphorylation kinetics of wild-type MbTEC^FL^ with MbTEC^FL^ bearing a mutated PRR motif (MbTEC^FL^ Gly_6_) (Figure 5F). We observed that MbTEC^FL^ Gly_6_ autophosphorylated considerably slower than MbTEC^FL^, suggesting that the polyproline motif plays an active role in MbTEC activation. A similar decrease in autophosphorylation was previously reported for human BTK mutated in its PRR (26). As observed previously, mutation of L396 in the SH2-kinase linker activated MbTEC, but addition of the PRR mutation (MbTEC^FL^ Gly_6_ L396A) did not have any additive effect. This finding is consistent with the situation in which the polyproline motif can compete with the SH2-kinase linker for binding to the SH3 domain under conditions in which autoinhibition by the PH domain is relieved.

In summary, activation of MbTEC is explicitly dependent on PIP_3_, which triggers displacement of the PH domain and subsequent engagement of the PRR by the SH3 domain, leading to relief of autoinhibition, autophosphorylation, and downstream substrate phosphorylation.

## Discussion

This study describes the activation of a TEC kinase by the lipid second messenger PIP_3_ with unprecedented mechanistic insight. Using a custom *in silico* pipeline for structure prediction with AlphaFold 3, we describe the nature of the intramolecular interface between the PIP_3_-binding PH domain and the kinase domain at atomic resolution. We validate the model with in-solution X-ray scattering data and HDX-MS analysis of truncation and interface mutants, and demonstrate a specific, physical interaction of the PH and kinase domains in *trans*. Finally, we present activity data supporting a model in which PIP_3_ elicits the allosteric activation of MbTEC by destabilizing an electrostatic network of inhibitory interactions between the PH and kinase domains and that this network serves to inactivate MbTEC in the absence of PIP_3_. Our findings converge on a model for the activation of MbTEC by PIP_3_ that is likely shared by all TEC kinases.

Our work has important implications for the accurate structure prediction of proteins and protein complexes. AlphaFold exploits deep co-evolutionary variance to predict protein structure. At its core is a MSA typically generated by profile hidden Markov models that can detect remote homologs. While this is a robust tool for predicting the fold of individual domains, it risks contaminating the MSA with partially homologous, but divergent sequences that mask evolutionary couplings between residues. Greater emphasis, therefore, needs to be placed on MSA curation to ensure that the input sequences belong to genuine protein orthologs. Similarly, our work emphasizes the risk of taxonomic bias in the MSA, something which AlphaFold does not explicitly address. Taxonomic bias arises from the over-representation of highly similar sequences and can lead to weaker signals for co-evolutionary variance. By addressing both of these issues, we were able to confidently predict the structure of full-length MbTEC where the standard AlphaFold implementation fails. While the importance of MSA curation, clustering, sampling and taxonomic balancing have all been previously reported (42–46), they have yet to be more widely adopted by the vast community of scientists employing AlphaFold in their studies.

A clear extension of these insights is that proper curation of an MSA will likely give rise to structural models that have a deep evolutionary conservation. In this case, the convergence on a singular model for the autoinhibited conformation of MbTEC implies that this conformation is conserved among all the TEC kinases. Further experimentation will no doubt be required to confirm this assumption, but literature evidence suggests that the model is compatible with the solution conformations of human ITK (21) and BTK revealed by NMR and HDX-MS respectively (20), meaning that we have already spanned the approximately 600-800 My of evolution between choanoflagellates and humans.

All of this leads us to propose a universal model for the activation of the TEC kinases by PIP_3_. Activation of cell surface receptors drives phosphorylation of their cytoplasmic tails by non-receptor tyrosine kinases (Figure 6). These phospho-tyrosine marks serve to recruit both the TEC kinase and class I PI3K. Recruitment of TEC alone is insufficient to drive its activation, which depends on the production of PIP_3_ by PI3K. Co-localization of autoinhibited TEC and PI3K at the plasma membrane increases the probability that TEC will be activated by PIP_3_ upon its production; indeed, XLA-associated mutations of the SH2 domain of BTK, including the phospho-tyrosine coordinating residue, R307, inhibit binding to the adaptor protein B-cell linker protein (BLNK) (47, 48) and impair BTK activation in cells (25). The SH2 domain of ITK is similarly required for its activation downstream of the TCR (49). Whilst PI(4,5,)P_2_ can bind to the PIP_3_ binding pocket of the PH domain, it is the specific recognition of the 3’ phosphate by residues K12, R28 and K53 that weakens the network of hydrogen bonds which stabilizes the autoinhibitory PH-kinase domain interface. Although we cannot exclude a model in which the interface dissociates prior to PIP_3_ binding, docking of PIP_3_ to the autoinhibited conformation of MbTEC is entirely compatible with molecular dynamics simulations of BTK that have revealed the orientation of the PH domain when bound to membrane-embedded PIP_3_ (19). Given that PIP_3_ is generated, and remains, in the plasma membrane, and that phosphorylated tyrosine motifs are locally constrained in the cytoplasmic tails of activated receptors, this spatially confines MbTEC signaling to the plasma membrane and temporally confines it to the lifetime of receptor activation. PIP_3_-mediated activation of TEC, coincident with the intramolecular engagement of its SH3 domain with its own polyproline motif, promotes its activation by autophosphorylation, an event that has been proposed to be mediated in *trans* by activation loop exchange (7). Upon attenuation of the PIP_3_ signal, dissociation of its PH domain from the membrane restores TEC autoinhibition, leading to immediate shutdown of the downstream signaling pathway.

**Figure 6.**
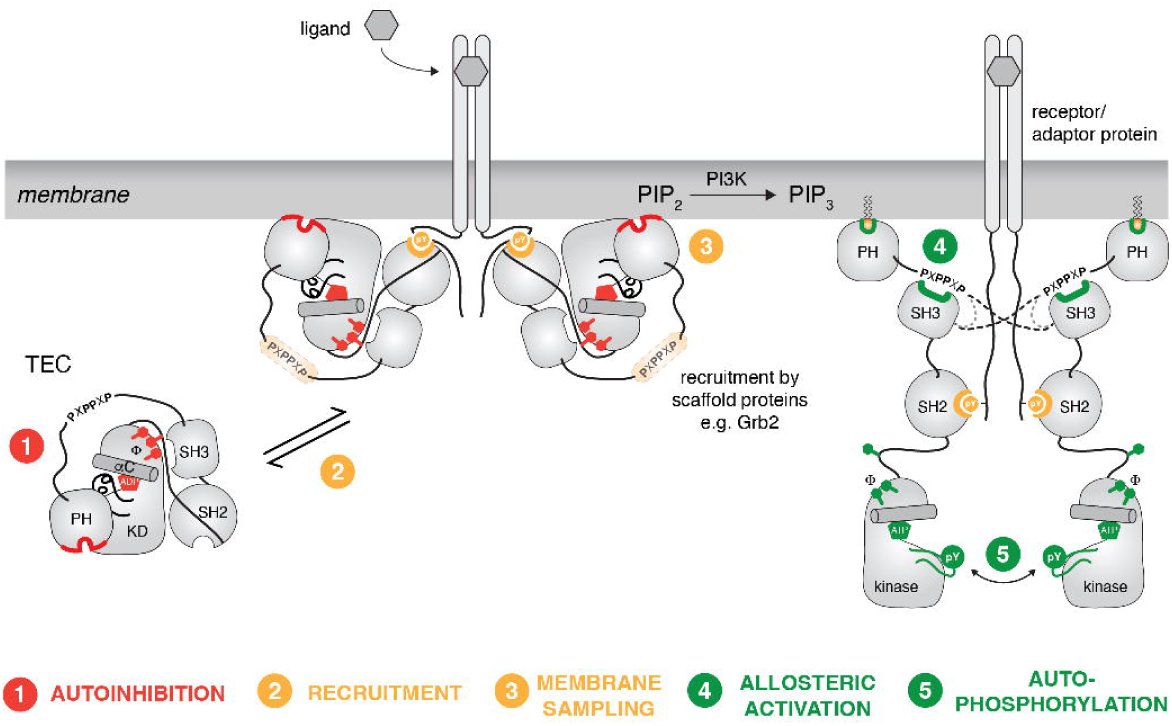
Model of PIP_3_-mediated auto-activation of TEC family kinases. TEC kinases adopt an autoinhibited conformation in the absence of an activating stimulus that is characterized by a discrete inhibitory interface between its PH and kinase domains (1). Upon ligand-mediated receptor activation and subsequent phosphorylation by non-RTKs, TEC is recruited to the membrane by phospho-tyrosine motifs in the receptor tails and SH3 domain-containing adaptor proteins such as Grb2 (2). Local concentration of TEC beneath the plasma membrane permits sampling of the membrane for PIP_3_ (3), engagement of which is concomitant with disengagement of its PH and SH3 domains from its kinase domain. PIP_3_ binding anchors TEC on the membrane and licenses its SH3 domain to compete with Grb2 for the polyproline motif in the PH-SH3 linker, thereby promoting allosteric activation of the kinase domain (4). Dimerization of TEC, driven by its local concentration on the membrane and possibly by reciprocal binding of the SH3 domain to the polyproline motif of a second TEC protomer, promotes activation loop autophosphorylation (5), thereby rendering TEC fully competent for downstream signaling. Upon loss of the PIP_3_ signal, TEC is autoinhibited by its PH domain, thereby eliciting an acute termination of downstream signaling.

Our finding that the PH domain is sufficient to attenuate >90% of kinase activity in the background of a mutation that mimics allosteric activation of the SH3-SH2-kinase module suggests that PIP_3_ can be considered the master regulator of TEC kinase activity. The coincident detection of each signal in space and time ensures the propagation of the signal with high fidelity and almost certainly limits the available substrate pool to those substrates co-localized to the site of TEC activation on the membrane. This is consistent with the membrane localization of one of the best characterized substrates of BTK, phospholipase C (PLC) γ (50, 51).

It would be remiss not to acknowledge the discrepancy between our structural model for MbTEC and experimental structures of human BTK that have been described by others (6, 7). Published SAXS data (7, 24, 25) and cryo-electron microscopy (7) on human BTK indicate a much more extended structure than we observe for MbTEC, while crystal structures of full-length BTK or synthetic PH-kinase domain fusion constructs either exhibited no electron density for the PH domain (7) or divergent contacts with the kinase domain (6, 7). As such, the prevailing model for autoinhibition by the PH domain involves an ensemble of different conformational states. However, caution must be exercised in the interpretation of published crystal structures, which may reflect the influence of lattice packing interactions in which enthalpically favorable interactions compensate for the reduction in entropy upon lattice formation. In the case of the solution scattering of BTK, we suspect that the scattering may reflect partial dimerization of BTK at the high concentrations required for SAXS. PH domain-mediated dimerization of BTK via insertion loops specific to BTK (Supplementary Figure 8A) has been extensively characterized and is supported by crystal structures of the so-called Saraste dimer (16, 17) (Supplementary Figure 8B). Here, it should be noted that PIP_3_-dependent dimerization of the PH domain of BTK, but not those of TEC or ITK, is required for BTK autophosphorylation and downstream signaling in B-cells (6, 14, 19, 52), indicating that paralog-specific evolutionary adaptations likely created additional layers of control over dimerization, kinase activation and, ultimately, signaling specificity.

Finally, whilst this study does not address the role of adaptor proteins in TEC kinase signaling, it offers new perspectives on their functions. Although our data is insufficient to exclude the possibility that adaptor proteins such as Grb2 directly activate TEC kinases (53) in cells, the high local concentration of the PRR and SH3 domain (intra-TEC) favors a TEC-autonomous model (Figure 6). Indeed, the *cis*-interaction of the polyproline motif in ITK has been shown to prevent the binding of SH3 domain-containing adaptor proteins such as Sam68 and Grb2 (39). In this sense, it is possible that adaptor proteins, such as Grb2, govern the subcellular localization of autoinhibited TEC kinases, whose activation is then accomplished in a PI3K-dependent and TEC-intrinsic manner. Such a model is consistent with the presence of a Grb2 ortholog in choanoflagellates (54) and would provide a coincidence-based mechanism for the discrete subcellular localization of TEC kinases with PI3K prior to their activation.

In conclusion, the TEC kinases are exquisitely tuned detectors that function in much the same way that an AND logic gate works in an electronic circuit. However, more sophisticated reconstitutions with interaction partners and substrates will undoubtedly be required to elucidate precisely what is necessary and sufficient for high-fidelity signaling.

## Supporting information

HDX-MS source data

## Acknowledgements

We thank Dr. Adam Round, who assisted in SAXS data collection on beamline BM29 at the ESRF, Grenoble. A plasmid encoding the DNA sequence of MbTEC and an expression plasmid for YopH were kind gifts from Dr. Todd Miller (Stony Brook University, NY, USA). This work was supported by the following grants from the Austrian Science Fund to T.A.L.: P28135, P30584, P33066, and P36212, and an operating grant from the Cancer Research Society (operating grant 1459101) to JEB.

## Author Contributions

Proteins were expressed and purified by E.K., F.v.R. and R.R. T.A.L. and L.P. performed all *in silico* modeling and computational analyses. All samples for SAXS analysis were purified by F.v.R. SAXS data were collected and analyzed by T.A.L. Kinase assays were performed by E.K. and F.v.R. HDX-MS experiments were performed by H.G.N. and E.E.W, with data analysis by H.G.N., E.E.W., and J.E.B. Thermal stability assays were performed by E.K. and F.v.R. L.P. prepared PIP_3_-containing lipid nanodiscs and performed all fluorescence anisotropy experiments. T.A.L. conceived the project. E.K., R.R. and T.A.L. wrote the manuscript. T.A.L. and J.E.B. obtained funding for the work.

## Materials and Methods

### Fluorescence anisotropy

For fluorescence anisotropy measurements of affinity, the MbTEC^PH^ was labeled with maleimide-Atto488 (ATTO-TEC) as previously reported (55). Unreacted dye was removed by gel filtration and labeling was confirmed by mass spectrometry (Supplementary Figure 1N). MbTEC^32K^ constructs were serially diluted in 20 mM HEPES pH 7.4, 150 mM NaCl, 2% glycerol and 1 mM TCEP containing 200 nM of the labeled PH domain. Measurements were performed with a FS5 Spectrofluorometer (Edinburgh Instruments) equipped with a QS High Performance cuvette (Hellma). For each concentration, 10 Technical replicates were measured at a controlled temperature of 21 °C. Excitation and emission wavelengths were set to 501 nm and 523 nm, with a bandwidth of 5 nm for both and an exposure time of 1 s. Affinities were determined by fitting a one-site binding model in GraphPad Prism.

### Radiometric kinase assay

All radiometric kinase assays were performed using radiolabeled [γ^32^ P] ATP (Hartman Analytic) at room temperature in the same buffer (20 mM HEPES pH 7.4, 150 mM NaCl, 2 % glycerol, 1 mM TCEP, 1 mM ATP and 2 mM MgCl_2_ spiked with 1:50 [γ^32^ P] ATP) and a final protein concentration of 0.5 - 2 μM. After clarifying the dephosphorylated protein sample, it was incubated for the indicated duration. At each time point a 10 μL sample was collected and the reaction was quenched by adding 2.5 μL of 1 M EDTA and 3 μL 5x SDS Loading Dye. The samples were separated by SDS-PAGE to remove non-incorporated ATP. The gel was washed 3x 5 min with water, dried and exposed to a phosphor screen overnight. The screen was imaged with an Amersham Typhoon imager and the radioactive signal was quantified with ImageJ. The fraction of phosphorylated MbTEC protein was calculated from an internal standard, assuming the incorporation of one phosphate per molecule MbTEC (verified by intact mass spectrometry).

For kinase assays containing PIP_3_ nanodiscs, MbTEC was incubated with nanodiscs in a 4-fold excess of protein (assuming a maximum binding stoichiometry of 4:1 MbTEC:nanodisc) for 1 h at room temperature prior to the addition of ATP and MgCl_2_. The final protein concentration was 1 μM.

### ADP-Glo kinase assay

Kinase assays were performed using the ADP-Glo kit (Promega) according to the manufacturer’s instructions. Kinase reactions contained 500 nM purified kinase, 50 μM SRC substrate (SignalChem #S30-58), and 1 mM ATP in 20 mM Tris pH 7.5, 100 mM NaCl, 1 mM TCEP, 2 mM MgCl_2_, and 0.05 mg/ml BSA. Reactions were incubated at RT for the specified time and luminescence was measured using a TECAN Infinite F500 plate reader.

### Small-angle X-ray scattering (SAXS)

SAXS data for MbTEC^FL^ and MbTEC^32K^ were collected on BM29 at the ESRF, Grenoble, France using an in-line SEC-SAXS setup. Proteins were applied to a Agilent Bio SEC 300 column equilibrated in 20 mM Tris, pH 7.4, 150 mM NaCl, 1 mM TCEP and 1 % (v/v) glycerol and images were acquired every second for the duration of the size exclusion run. Buffer subtraction was performed by averaging 50 frames either side of the peak. All subsequent data processing steps were performed using the ATSAS data analysis software 3.9.1. The program DATGNOM (56) was used to generate the pair distribution function [P(r)] for each isoform and to determine D_max_ and R_g_ from the scattering curves [I(q) vs. q] in an automatic, unbiased manner. For *ab initio* calculation of the molecular envelopes, iterative rounds of dummy atom modeling were followed by alignment of the resulting models, trimming of outliers, and refinement of the core electron density, followed by superposition of the respective models for MbTEC^FL^ and MbTEC^32K^.

