## Supplementary Information for "Mechanism of activation of an ancestral TEC kinase by PIP_3_"

Contains: Supplementary Figures 1-8
Supplementary Methods
Supplementary References

### Supplementary Figures

#### Supplementary Figure 1. Mass spectrometry of constructs employed in this study.

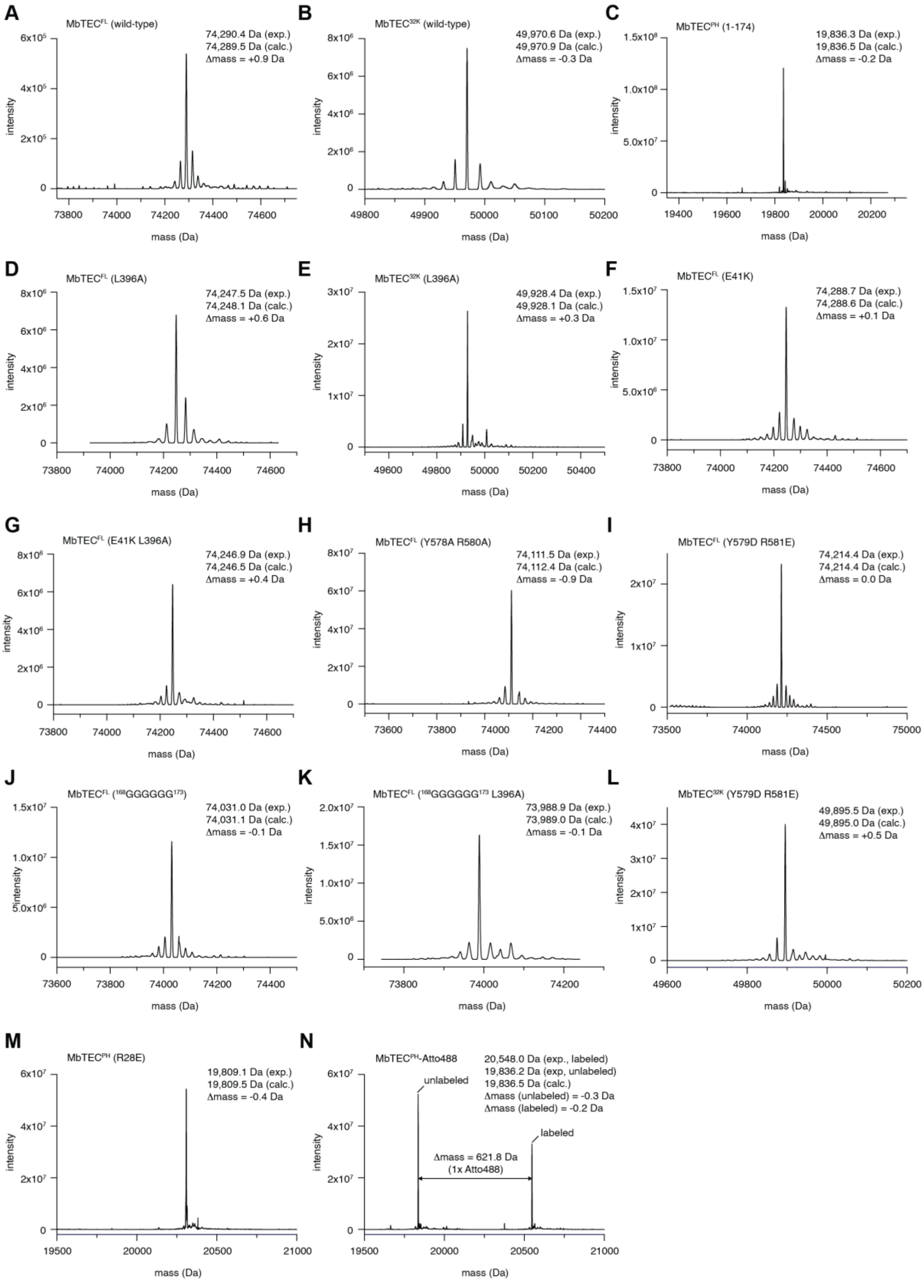

A. Deconvoluted mass spectrum of MbTEC$^{FL}$.

B. Deconvoluted mass spectrum of MbTEC$^{32K}$.

C. Deconvoluted mass spectrum of MbTEC$^{PH}$.

D. Deconvoluted mass spectrum of MbTEC$^{FL}$ L396A.

E. Deconvoluted mass spectrum of MbTEC$^{32K}$ L396A.

F. Deconvoluted mass spectrum of MbTEC$^{FL}$ E41K.

G. Deconvoluted mass spectrum of MbTEC$^{FL}$ E41K L396A.

H. Deconvoluted mass spectrum of MbTEC$^{FL}$ Y579A R581A.

I. Deconvoluted mass spectrum of MbTEC$^{FL}$ Y579D R581E.

J. Deconvoluted mass spectrum of MbTEC$^{FL}$ $^{168}$GGGGGG$^{173}$.

K. Deconvoluted mass spectrum of MbTEC$^{FL}$ $^{168}$GGGGGG$^{173}$ L396A.

L. Deconvoluted mass spectrum of MbTEC$^{32K}$ Y579D R581E.

M. Deconvoluted mass spectrum of MbTEC$^{PH}$ R28E.

N. Deconvoluted mass spectrum of MbTEC$^{PH}$-Atto488.

Supplementary Figure 2. Biophysical characterization of constructs employed in this study.

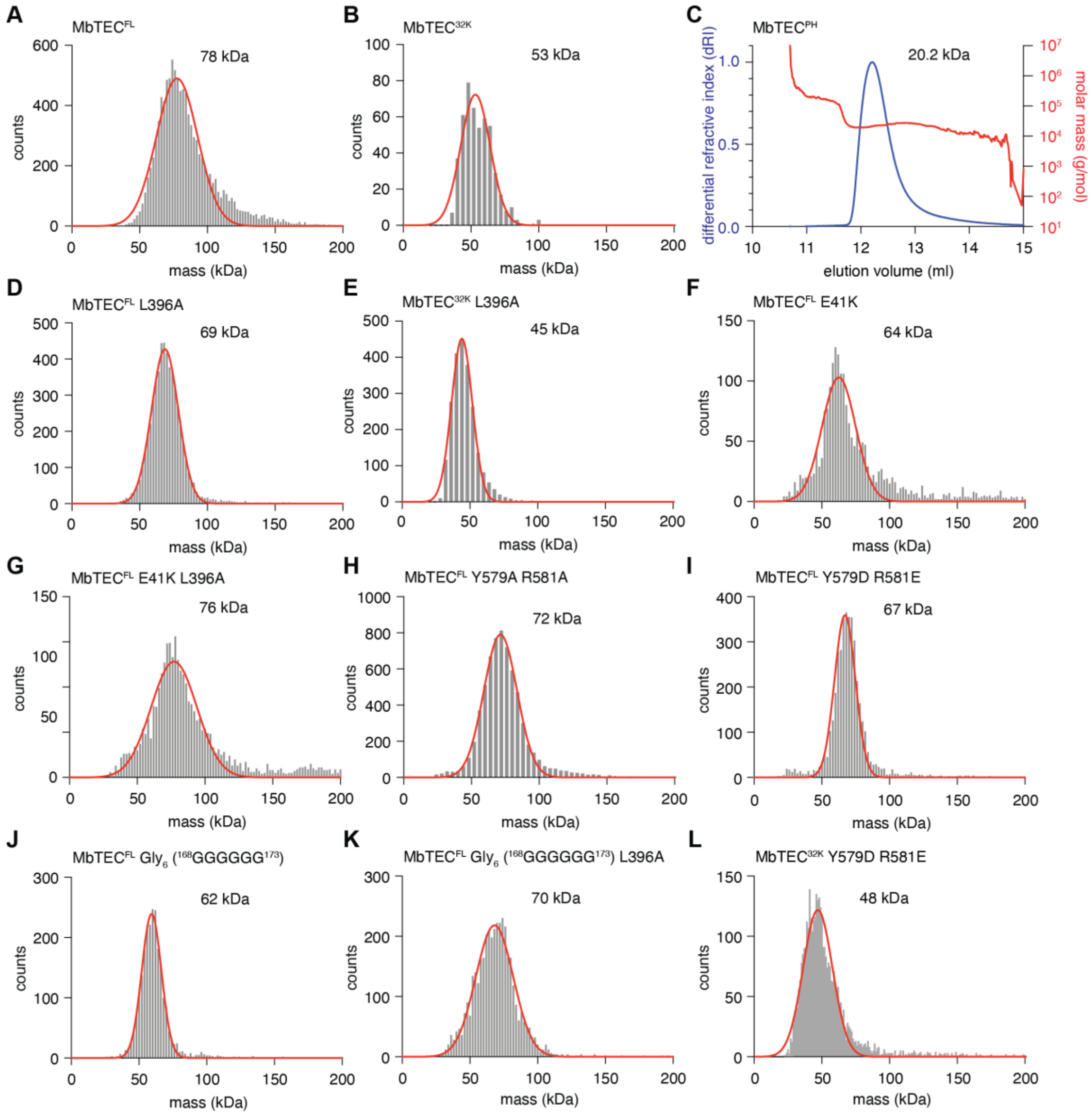


A. Mass photometry of MbTEC^FL.

B. Mass photometry of MbTEC^32K.

C. SEC-MALS analysis of MbTEC^PH.

D. Mass photometry of MbTEC^FL L396A.

E. Mass photometry of MbTEC^32K L396A.

F. Mass photometry of MbTEC^FL E41K.

G. Mass photometry of MbTEC$^{FL}$ E41K L396A.

H. Mass photometry of MbTEC$^{FL}$ Y579A R581A.

I. Mass photometry of MbTEC$^{FL}$ Y579D R581E.

J. Mass photometry of MbTEC$^{FL}$ Gly6 ($^{168}$GGGGGG$^{173}$).

K. Mass photometry of MbTEC$^{FL}$ Gly6 ($^{168}$GGGGGG$^{173}$) L396A.

L. Mass photometry of MbTEC$^{32K}$ Y579D R581E.

Supplementary Figure 3. *M. brevicollis* TEC adopts a conserved autoinhibited conformation.

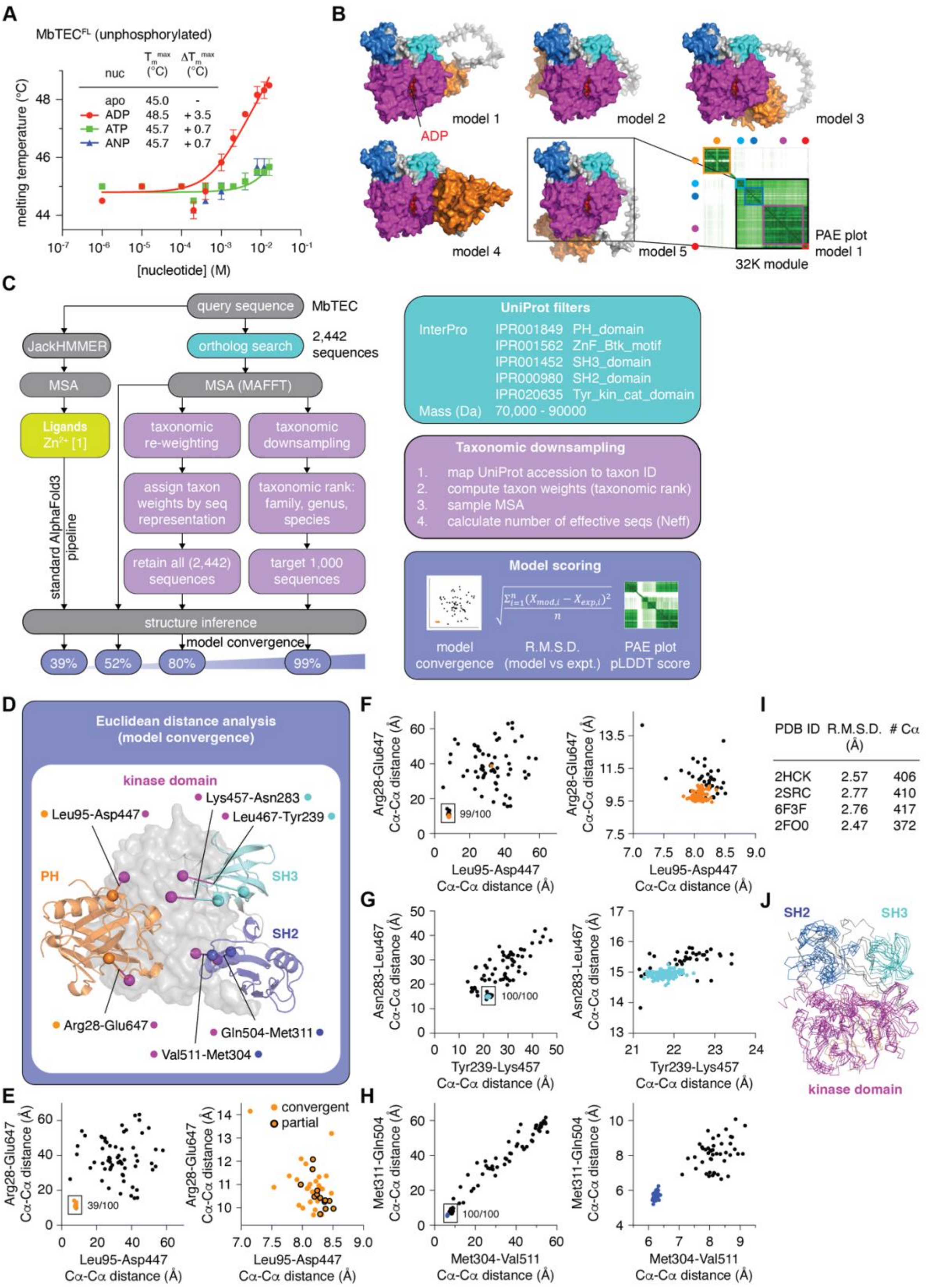

A. Thermal stability measurements of MbTec as a function of nucleotide concentration. ADP (red), ATP (green), AMPPNP (ANP; blue).

B. Five independent AlphaFold 3 predictions of the structure of human Btk. Bottom right: pair alignment error (PAE) plot for model 1.

C. Pipeline for custom MSA building, curation, and taxonomic reweighting or downsampling. The custom MSA is then fed into the structure inference step of AlphaFold 3 along with ligands (in the case of MbTEC, a single zinc ion). We scored the output models using three metrics: Euclidean distance plots (convergence on domain arrangements), root mean square deviation (R.M.S.D.) of the model from experimental structures, the PAE plot (pair-aligned residue model confidence). Improvement of custom AlphaFold 3 structure prediction pipelines on model convergence compared to the standard configuration of AlphaFold 3 implemented on the publicly accessible AlphaFold 3 server.

D. Schematic of model evaluation by Euclidean distance measurements for MbTEC. The distance between fixed residue pairs on the kinase domain and each other regulatory domain (PH, SH3, SH2) served as a proxy for the relative domain arrangements in each model. The residue pairs are indicated with colored spheres joined by lines. The color of the sphere indicates the domain to which the residue belongs.

E. Outcome of 100 independent predictions using the AlphaFold 3 server. Input: MbTEC sequence and a single zinc ion. 39/100 models (orange circles) converged on the same arrangement of the PH and kinase domains, of which 26 also converged on the same arrangement of the SH3, SH2 and kinase domains (convergent models). The remaining 13 models (partial convergence) exhibited divergent positions for the SH3 and SH2 domains with respect to the PH and kinase domains (orange circles, black outline).

F. PH-kinase domain Euclidean distance analysis of 100 independent AlphaFold 3 models using the standard pipeline (black circles) compared to our custom pipeline (orange circles).

G. SH3-kinase domain Euclidean distance analysis of 100 independent AlphaFold 3 models using the standard pipeline (black circles) compared to our custom pipeline (cyan circles).

H. SH2-kinase domain Euclidean distance analysis of 100 independent AlphaFold 3 models using the standard pipeline (black circles) compared to our custom pipeline (blue circles).

I. Table of PDB entries for experimentally determined structures of the 32K module of TEC kinases and their corresponding R.M.S.D. from the highest ranked AlphaFold 3 model obtained (model 18).

J. Superposition of the MbTEC32K model with the corresponding structures of Src (2SRC, 6F3F), Hck (2HCK), and Abl (2FO0).

Supplementary Figure 4. MbTEC adopts a compact conformation in solution.

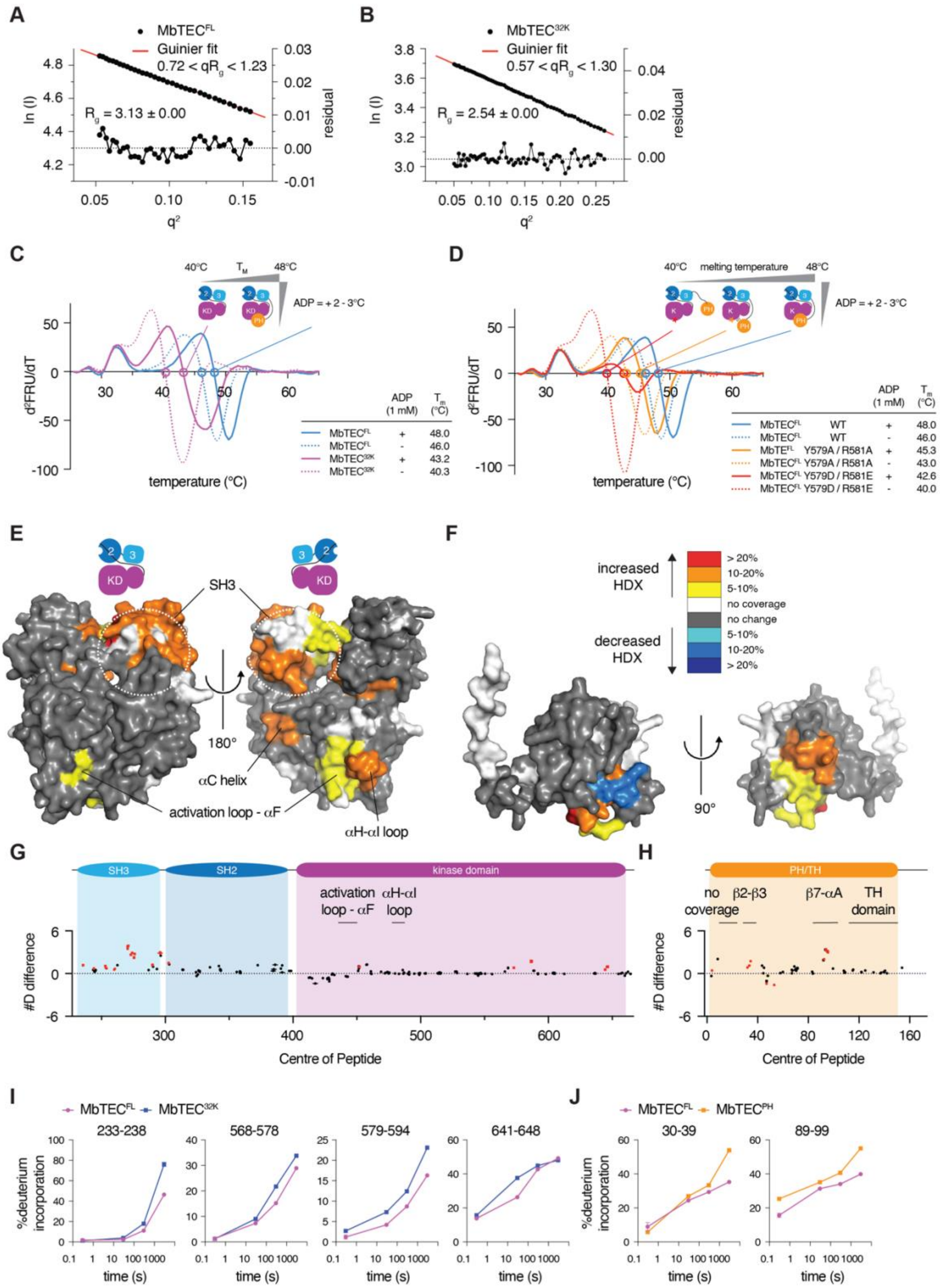

A. Guinier fit of the low-angle scattering regime for MbTEC$^{FL}$.

B. Guinier fit of the low-angle scattering regime for MbTEC$^{32K}$.

C. Second derivatives of the melting curves for $MbTEC^{FL}$ (blue), $MbTEC^{32K}$ (magenta). Solid lines: + 1mM ADP; dotted lines: -ADP. Inset: table of corresponding MbTEC melting temperatures in the presence and absence of 1 mM ADP and 2 mM $MgCl_2$.

D. Second derivatives of the melting curves for $MbTEC^{FL}$ (blue), $MbTEC^{FL}$ Y579A R581A (orange) and $MbTEC^{FL}$ Y579D R581E (red). Solid lines: + 1mM ADP; dotted lines: -ADP. Inset: table of corresponding MbTEC melting temperatures in the presence and absence of 1 mM ADP and 2 mM $MgCl_2$.

E. Pairwise HDX-MS analysis of $MbTEC^{FL}$ compared to $MbTEC^{32K}$. Significant differences in HDX in $MbTEC^{32K}$ are mapped onto the AlphaFold3 model of $MbTEC^{32K}$ and color coded according to the legend.

F. Pairwise HDX-MS analysis of $MbTEC^{FL}$ compared to $MbTEC^{PH}$. Significant differences in HDX in $MbTEC^{PH}$ are mapped onto the AlphaFold3 model of $MbTEC^{PH}$ and color coded according to the legend.

G. Sum of #D difference in deuterium incorporation for $MbTEC^{32K}$ over the entire time course. Each point represents and individual peptide with significant changes shown in red (greater than 0.40 Da, 5% difference and two tailed t-test $p<0.01$ at any timepoint, n=3). Domain architecture is annotated above.

H. Sum of #D difference in deuterium incorporation for $MbTEC^{PH}$ over the entire time course. Each point represents an individual peptide with significant changes shown in red (greater than 0.40 Da, 5% difference and two tailed t-test $p<0.01$ at any timepoint, n=3). Domain architecture is annotated above. The full deuterium exchange information for all peptides is included in the source data.

I. Deuterium incorporation plots for selected significant peptides shown in panel G.

J. Deuterium incorporation plots for selected significant peptides shown in panel H.

#### Supplementary Figure 5. MbTEC is autoinhibited by its PH domain.

**A**

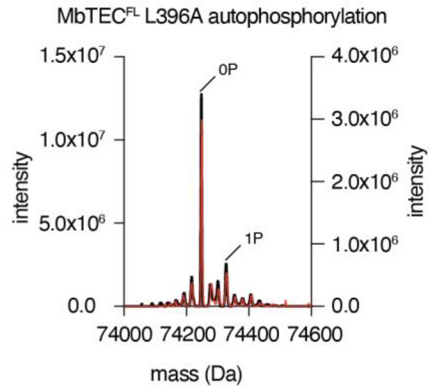


**B**

MS/MS of autophosphorylated MbTEC L396A (trypsin digest)

MbTEC$^{FL}$ L396A coverage = 85.2%
MbTEC$^{32K}$ L396A coverage = 79.4%

| Position | Sequence | ratio mod/base MbTEC$^{32K}$ L396A | ratio mod/base MbTEC$^{FL}$ L396A |
|---|---|---|---|
| 559 | FVIDDEYTASEGTK (2) | 0.03 | 0.00 |
| 559 | FVIDDEYTASEGTKFPIK (2) | 0.30 | 0.02 |
| 559 | FVIDDEYTASEGTKFPIK (3) | 0.42 | 0.04 |

A. Mass spectrometry analysis of the product of MbTEC$^{FL}$ L396A autophosphorylation (denoted by the black circle in panel A). Intensities before autophosphorylation in black (left axis) and after autophosphorylation in red (right axis).

B. Tandem mass spectrometry identification of the modified residues in MbTEC$^{32K}$ L396A.

Supplementary Figure 6. Membrane translocation of MbTEC$^{FL}$ and MbTEC$^{PH}$ in cells.

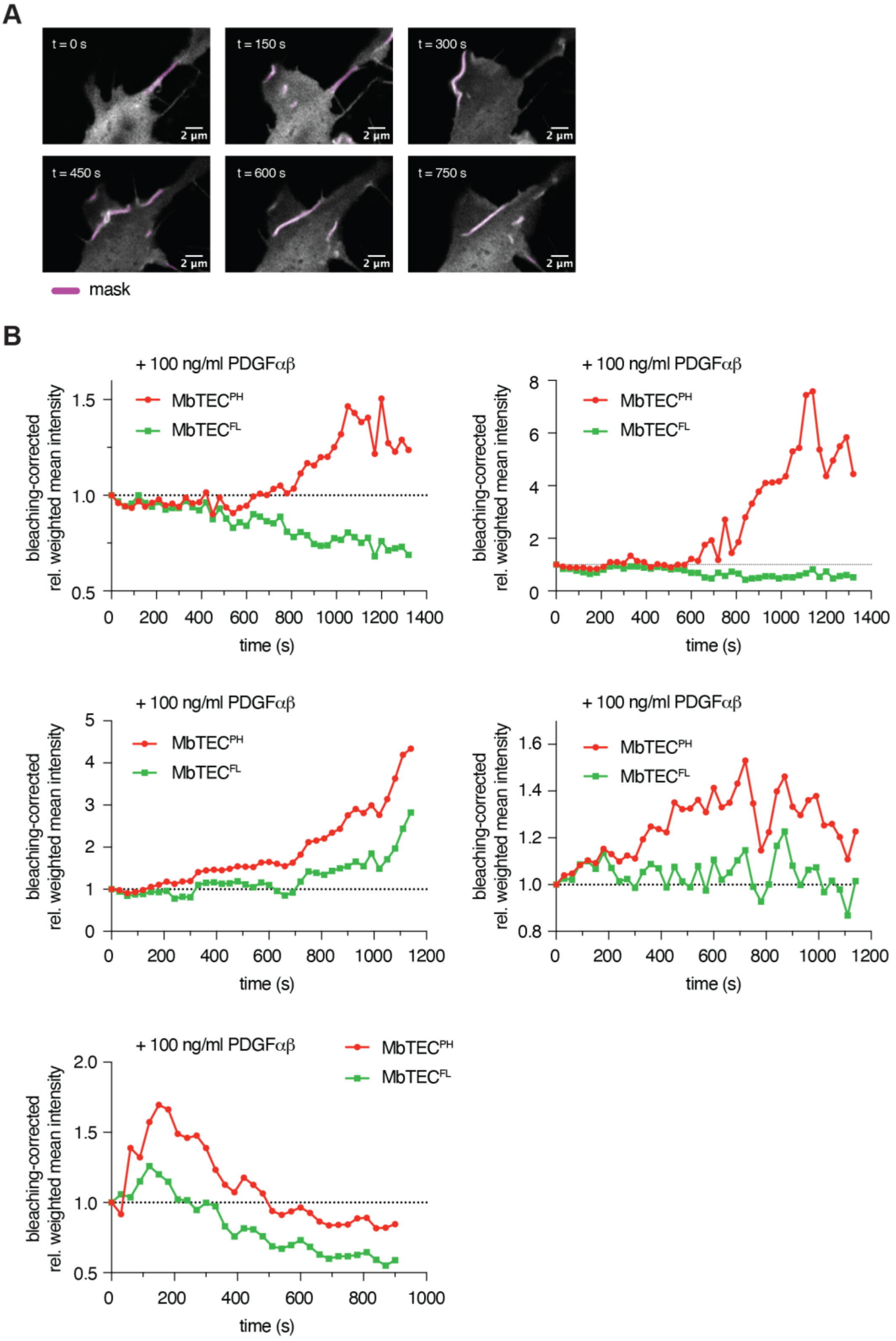


A. Plasma membrane association of EGFP-MbTEC$^{FL}$ and mCherry-MbTEC$^{PH}$ in a representative NIH3T3 fibroblast following stimulation with PDGFαβ. Mask of filopodia-like membrane structures used for signal integration shown in magenta.

B. Bleaching-corrected, normalized mask intensity profiles of $MbTEC^{FL}$ (green) and $MbTEC^{PH}$ (red) in five independent cells post-PDGF$\alpha\beta$ stimulation as a function of time.

#### Supplementary Figure 7. MbTEC is allosterically activated by $PIP_3$.

**A**

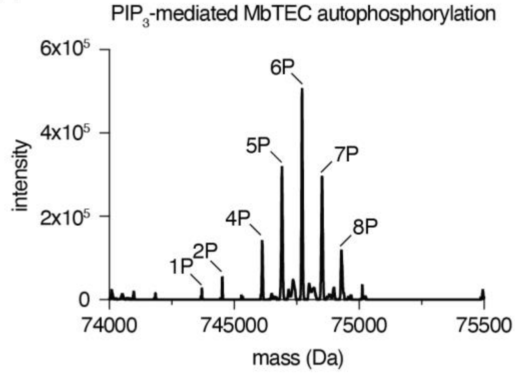


**B**

*MbTEC*

| Position | Sequence window | Probability | Modification multiplicity | # sequences | # modified sequences | Spectral count |
|---|---|---|---|---|---|---|
| 279 | AGQEGYIPANY | 1.00 | 1;2 | 5 | 9 | 157 |
| 284 | YIPANYVRKLG | 1.00 | 1;2 | 5 | 9 | 131 |
| 336 | SLSVFYQGKLK | 1.00 | 1;2 | 3 | 6 | 10 |
| 352 | HDQDMYRVSER | 1.00 | 1 | 4 | 6 | 22 |
| 369 | SELIEYHKHNG | 1.00 | 1 | 2 | 2 | 11 |
| 427 | VRAGVYKSNRP | 1.00 | 1 | 2 | 2 | 16 |
| 559 | VIDDEYTASEG | 1.00 | 1 | 2 | 2 | 15 |

**C**

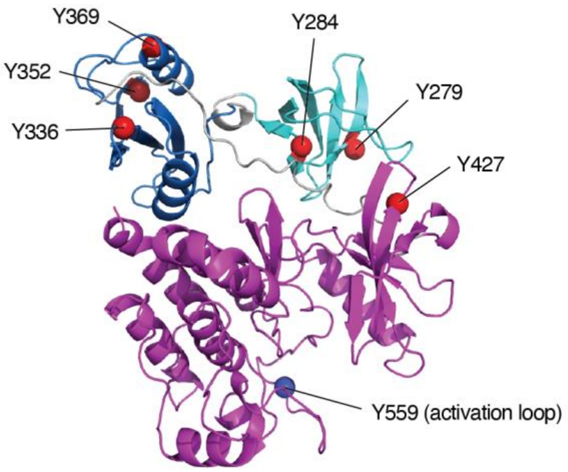


**D**

*Membrane scaffold protein 1 (MSP1D1)*

| Position | Sequence window | Probability | Modification multiplicity | # sequences | # modified sequences | Spectral count |
|---|---|---|---|---|---|---|
| 47 | AKVQPYLDDFQ | 0.99 | 1 | 1 | 1 | 2 |
| 62 | EEMELYRQKVE | 1.00 | 1 | 5 | 5 | 7 |
| 113 | THLAPYSDELR | 1.00 | 1 | 4 | 4 | 9 |
| 139 | ARLAEYHAKAT | 1.00 | 1 | 3 | 3 | 8 |
| 183 | SALEEYTKKLN | 1.00 | 1 | 2 | 2 | 2 |

A. Intact mass spectrometry of MbTEC after 90 min autophosphorylation on $PIP_3$-containing nanodiscs.

B. Tandem mass spectrometry analysis of autophosphorylated MbTEC following activation by $PIP_3$ nanodiscs.

C. Phospho-sites identified in panel A mapped onto predicted structure of $MbTEC^{32K}$.

D. Phospho-sites in MSP1D1 identified by tandem mass spectrometry.

#### Supplementary Figure 8. Paralog-specific differences in the PH domain of Tec kinases.

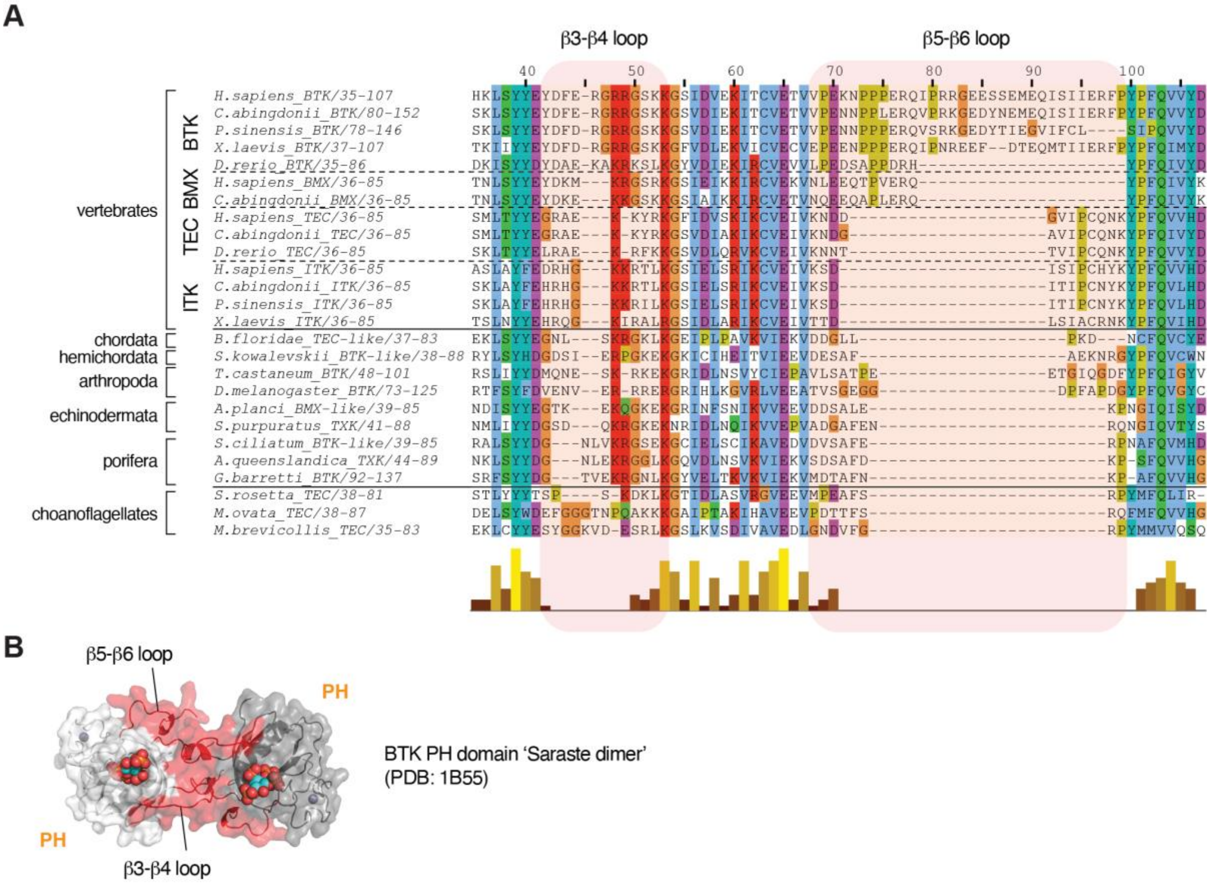


A. Partial multiple sequence alignment of the PH domain of the TEC kinases over 600-800 My of evolution. Loop insertions β3-β4 and β5-β6 specific to BTK are highlighted with a red background.

B. Crystal structure of the 'Saraste' dimer of the BTK PH domain, indicating that the β3-β4 and β5-β6 insertion loops in BTK mediate dimerization.

### Supplementary Methods

#### Structure prediction using AlphaFold 3

A curated list of 2,442 MbTEC homologs was generated by filtering UniProt for sequences containing the following domains, as defined by InterPro: PH (IPR001859), Btk_Zn_finger (IPR001562), SH3 (IPR001452), SH2 (IPR000980), and tyrosine kinase (IPR020635). A mass range of 70,000-90,000 Da was included to filter out sequences of excessive length. The sequences were aligned using MAFFT (1, 2) with default settings (BLOSUM62 scoring matrix, gap opening penalty: 1.53, offset value: 0.0) and converted into the A3M format required by AlphaFold. The custom MSA was either fed directly into the database search step of AlphaFold 3 (3) or further curated by taxonomic reweighting or downsampling. For taxonomic reweighting, individual sequences in the MSA were reweighted according to their taxonomic representation such that individual sequences in over-represented taxa were downweighted for the structure inference step. All 2,442 sequences were retained in the reweighted MSA.

##### Taxonomic downsampling

To reduce bias from over-represented taxa while preserving evolutionary diversity, the alignment was taxonomically downsampled at the *family* level using a custom Python pipeline. Each sequence $i$ was assigned a raw weight

$$w_i^{\text{raw}} = \frac{1}{n_{\text{family}(i)}},$$

where $n_{\text{family}(i)}$ is the total number of sequences from the same taxonomic family as sequence $i$. The raw weights were normalized with

$$w_i = \frac{w_i^{raw}}{\sum_j w_j^{raw}}$$

so that all weights sum to 1. Sequences from different families were randomly sampled proportionally to the normalized weights until reaching a target alignment size (1000 sequences). The effective number of sequences ($N_{\text{eff}}$) in the MSA was calculated from the sampling weights as

$$N_{\text{eff}} = \frac{\sum_i w_i}{max_i w_i},$$

which, for normalized weights, reduces to $N_{\text{eff}} = 1/\max_i w_i$. $N_{\text{eff}}$ reflects the diversity of the weighted alignment and is commonly used as a proxy for the amount of independent evolutionary information available for co-evolutionary and structure-prediction methods. The procedure and the definition of $N_{\text{eff}}$ follow those used in previous work on evolutionary-coupling-based structure prediction (4, 5). Calculation of $N_{\text{eff}}$ before and after taxonomic downsampling revealed a reduction from 719 to 295, representing fewer but more diverse sequences. The custom MSA (2,442 sequences, no reweighting or downsampling), reweighted MSA (2,442 sequences, reweighted by taxonomic family), or balanced MSA (1,000 sequences, downsampled by taxonomic family) MSAs were fed into the structure inference pipeline of AlphaFold 3.

##### Structure inference

For each curated MSA, 100 models were predicted from random seeds. A seed in AlphaFold 3 represents a random starting pose, from which the final model is obtained by diffusion (3). Model convergence was assessed by calculating the Euclidean distances of residues on the kinase domain to residues on the PH, SH3 and SH2 regulatory domains (Supplementary Figure 3E), using the formula $d_{1,2} = \sqrt{(x_1 - x_2)^2 + (y_1 - y_2)^2 + (z_1 - z_2)^2}$ with x, y, and z being the residue Cα coordinates. The agreement with experimental data was assessed by calculation of the root mean square deviation over corresponding main chain Cα atoms for known structures of the SH3-SH2-KD (32K) module. Finally, the models were evaluated by AlphaFold ranking score and the pair alignment error (PAE) plot for the best scoring model.

#### Protein expression and purification

The sequence for M. brevicollis MX1 TEC was obtained from MONBRscaffold_8 20778 to 23025 (-). $MbTEC^{FL}$ (1-667) and $MbTEC^{32K}$ (229-667) were cloned into the pFastBac Dual vector with an N-terminal GST tag followed by a TEV cleavage site for expression in baculovirus-infected Sf9 insect cells. MbTEC mutants were generated by site-directed mutagenesis.

Cells were centrifuged at 4000 g for 30 min and the pellet lysed in 100 mL lysis buffer (50 mM HEPES pH 7.4, 150 mM NaCl, 1 mM TCEP, 0.25 % CHAPS, 2 mM $MgCl_2$, 1x protease inhibitor cocktail (Sigma P8849)) per 1 L expression culture. The lysate was centrifuged at 39000 g for 30 min at 4 °C and the supernatant was incubated with 5 mL glutathione Sepharose beads (Cytiva) for 2 h at 4°C. The beads were washed three times with Buffer A (50 mM Hepes pH 7.4, 50 mM NaCl, 1 mM TCEP, 0.25 % CHAPS), three times with Buffer B (50 mM Hepes pH 7.4, 300 mM NaCl, 1 mM TCEP, 0.25 % CHAPS) before being resuspended in Buffer C (50 mM Hepes pH 7.4, 150 mM NaCl, 1 mM TCEP, 0.25 % CHAPS). The protein was cleaved off the beads by adding 8.8 nmol TEV protease (purified in-house) and dephosphorylated by adding 10 nmol lambda phosphatase and 13 nmol YopH phosphatase (both purified in-house) overnight at 4 °C. Cleaved MbTEC protein was separated from the beads and diluted to 25 mM NaCl in Buffer $Q_A$ (50 mM HEPES, pH 7.4, 1 mM TCEP). The protein was loaded onto a MonoQ anion exchange column (Cytiva). After gradient elution with Buffer $Q_B$ (50 mM Hepes pH 7.4, 500 mM NaCl, 1 mM TCEP) from 0 to 50 % B in 520 column volumes (CV) to separate differentially phosphorylated protein species, the peak fractions were pooled, concentrated and injected onto a S200 10/300 GL size exclusion chromatography (SEC) column (Cytiva) equilibrated in SEC Buffer (20 mM HEPES pH 7.4, 150 mM NaCl, 2 % glycerol, 1 mM TCEP). The phosphorylation state of every construct was verified by intact mass spectrometry.

MbTEC$^{PH}$ (1-174) was cloned into the pGST parallel expression vector with an N-terminal GST tag preceded by a TEV cleavage site for expression in BL21 STAR bacterial cells. The cells were grown in LB medium containing 100 μg/mL ampicillin at 37 °C, 200 rpm to an OD = 0.7 before induction with 250 μM IPTG and incubation overnight at 20 °C. The cells were centrifuged at 4000 g for 30 min and the pellet lysed in 50 mL lysis buffer (50 mM HEPES pH 7.4, 150 mM NaCl, 1 mM TCEP, 0.25 % CHAPS, 2 mM $MgCl_2$, 0.2 mg/mL lysozyme, 2 mM benzamidine, 1 % glycerol) per 1 L expression culture. The lysate was sonicated, centrifuged at 39000 g for 30 min at 4 °C and the supernatant was incubated with 5 mL glutathione Sepharose beads (Cytiva) for 2 h at 4 °C. The beads were washed three times with Buffer A (50 mM Hepes pH 7.4, 50 mM NaCl, 1 mM TCEP, 0.25 % CHAPS), three times with Buffer B (50 mM Hepes pH 7.4, 300 mM NaCl, 1 mM TCEP, 0.25 % CHAPS) before being resuspended in Buffer C (50 mM Hepes pH 7.4, 150 mM NaCl, 1 mM TCEP, 0.25 % CHAPS). The protein was cleaved off the beads by adding

8.8 nmol TEV protease (purified in house) overnight at 4 °C. Cleaved $MbTEC^{PH}$ protein was separated from the beads and diluted to 30 mM NaCl in Buffer $S_A$ (50 mM HEPES, pH 7.4, 1 mM TCEP). The protein was loaded onto a CaptoS cation exchange column (Cytiva). After gradient elution with Buffer $S_B$ (50 mM HEPES pH 7.4, 500 mM NaCl, 1 mM TCEP) from 0 to 100 % B in 50 CV, the peak fractions were pooled, concentrated and injected onto a S75 10/300 GL size exclusion chromatography column (Cytiva) equilibrated in SEC Buffer (20 mM HEPES pH 7.4, 150 mM NaCl, 2 % Glycerol, 1 mM TCEP).

#### Thermal stability assays

Differential scanning fluorimetry measurements were performed in SEC buffer with 5x SYPRO Orange (life technologies) using a BioRad iQ5 Multicolor Real-Time PCR detection system. For each sample, the protein was diluted to 0.1 mg/mL either in the presence or absence of 1 mM ADP and 2 mM $MgCl_2$. The samples were heated from 25 °C to 90 °C in increments of 0.5 °C. The melting temperature was determined from the second derivative of the melting curve and averaged from three independent measurements.

#### Mass spectrometry

##### Intact mass spectrometry

Intact protein samples were diluted in $H_2O$ and up to 100 ng protein were loaded on an XBridge Protein BEH C4 column (2.5 µm particle size, dimensions 2.1 mm X 150 mm; Waters) using a Dionex Ultimate 3000 HPLC system (Thermo Fisher Scientific) with a working temperature of 50 °C, 0.1% formic acid (FA) as solvent A, 80% acetonitrile, 0.08% FA as solvent B. Proteins were separated with a 6 min step gradient from 10 to 80% solvent B at a flow rate of 300 µL/min and analysed on a Synapt G2-Si coupled via a ZSpray ESI source (Waters). Data were recorded with MassLynx V 4.1 (Waters) and analyzed using the MaxEnt 1 process to reconstruct the uncharged average protein mass.

#### Tandem mass spectrometry

##### *Sample preparation*

2 ug protein were precipitated overnight at -20°C by adding ice-cold acetone to a concentration of 80%. The proteins were pelleted by centrifugation at 16000 g for 10 min at 18°C. The pellets were dissolved in 2% sodium deoxycholate (SDC) in 50 mM ammonium bicarbonate (ABC). Proteins were reduced by adding 250 mM dithiothreitol (DTT) to final concentration of 10 mM for 30 min at 50°C and alkylated by adding 500 mM iodoacetamide (IAA) to final concentration of 20 mM and incubating for 30 min at room temperature in the dark. The remaining IAA was quenched for 10 min by adding 5 mM DTT. The samples were diluted to 1% SDC with 20 µL 50 mM ABC. Proteins were digested by adding 60 ng trypsin (Trypsin Gold, Promega; 100ng/µL solution in 1mM HCl) and incubated at 37°C overnight. The digest was stopped by the addition of 10% trifluoroacetic acid (TFA) to reach a final concentration of 2%. SDC was pelleted by centrifugation at 18,500 x g for 10 minutes. The resulting supernatant was desalted using C18 Stagetips (6).

##### *Liquid chromatography mass spectrometry*

LC-MS analysis was performed on a Vanquish Neo UHPLC system (Thermo Scientific) coupled to an Orbitrap Exploris 480 mass spectrometer (Thermo Scientific). The system was equipped with a Nanospray Flex ion source (Thermo Scientific), coated emitter tips (PepSep, MSWil), and a Butterfly Portfolio Heater (Phoenix S&T).

Peptides were loaded onto a trap column (Acclaim PepMap 100 C18 HPLC Column, 20 mm × 0.1 mm, 5 µm particle size, Thermo Scientific) using 0.1% TFA as mobile phase, and separated on an analytical column (Acclaim PepMap 100 C18 HPLC Column, 50 cm × 75 µm, 2 µm particle size, Thermo Scientific), applying a linear gradient starting with a mobile phase of 98% solvent A (0.1% FA) and 2% solvent B (80% acetonitrile, 0.08% FA), increasing to 35% solvent B over 60 min at a flow rate of 230 nl/min. The analytical column was heated to 30°C.

The mass spectrometer was operated in data-dependent acquisition (DDA) mode, with 2 s MS1 cycle time. Survey scans were acquired from 375-2000 m/z with lock mass enabled, normalized AGC target of 300%, resolution of 120,000. The most intense precursor ions (charge states +2 to +6) were selected for fragmentation using an isolation window of 1.4 m/z. Selected ions were analyzed with a maximum fill time of

200 ms, normalized AGC target of 200%, and resolution of 30,000 after HCD fragmentation with normalized collision energy of 30%. Monoisotopic precursor selection (MIPS) was set to “peptide” mode with relax restrictions on, the intensity threshold to 2.5 E4, and selected precursors were dynamically excluded for 20 seconds with isotope exclusion enabled.

##### *Mass spectrometry data analysis*

MS raw data were analyzed with FragPipe (22.0), using MSFragger (4.1) (7), IonQuant (1.10.27) (8), and Philosopher (5.1.1) (9). The default FragPipe workflow was used, turning on MS1 Quant omitting normalization and MBR. Cleavage specificity was set to Trypsin/P, with two missed cleavages allowed. The ion, peptide and protein FDR was set to 1%. Carbamidomethyl was used as fixed cysteine modification; methionine oxidation, protein N-terminal acetylation and phosphorylation of serine, threonine and tyrosine were specified as variable modifications. MS2 spectra were searched against the *S.frugiperda* entries from Uniprot (Taxon ID: 7108, release 2024.01), concatenated with a database of 379 common laboratory contaminants (release 2023.03, https://github.com/maxperutzlabs-ms/perutz-ms-contaminants), and the entries for the MbTEC constructs.

Computational analysis was performed using Python and the Python library MsReport (0.0.26) (10). Protein intensities were log2 transformed, phosphorylation sites with a localization probability < 80% were filtered out. The Python library XlsxReport (0.1.1) (11) was used to create formatted Excel files summarizing the results of the proteomics experiments. The proteomics data have been deposited to the ProteomeXchange Consortium via the PRIDE partner repository (12) with the dataset identifier PXD064005.

#### Mass photometry

Microscopy coverslips were cleaned by sequential sonication in water, isopropanol and water before drying under an air stream. Prior to measurements, CultureWell gaskets (Merck) were mounted onto the coverslip. Protein samples were clarified by centrifugation (5 min, 21000 g, 4 °C diluted to 0.5 μM in SEC buffer. To find focus, 19 μL of buffer were added into a chamber, the focal position was identified and secured with an autofocus system. Then, 1 μL of 0.5 μM protein sample was added, mixed and movies of 60 s duration were recorded. All measurements were performed using a Two$^{MP}$ mass photometer (Refeyn) and analyzed using the software Discover MP (Refeyn). Contrast-to-mass calibration was performed before every experiment.

#### Size exclusion chromatography coupled to multi-angle light scattering (SEC-MALS)

The oligomeric state of the PH domain construct was assessed by SEC-MALS. 50 µL of PH domain (3 mg/mL) was injected onto a Superdex 75 10/300 column (Cytiva) connected to a 1260 Infinity HPLC (Agilent Technologies) equilibrated in 50 mM HEPES pH 7.4, 150 mM NaCl, 1 mM TCEP. Light scattering was detected with a MiniDawn Treos (Wyatt) and the refractive index was measured with a Shodex RI-101 (Shodex) detector.

#### HDX-MS

##### Sample preparation

HDX reactions comparing MbTEC$^{FL}$ to either MbTEC$^{FL}$ Y579D R581E, MbTEC$^{32K}$ construct or MbTEC$^{PH}$ were carried out in a 20 µl reaction volume containing 15 pmol (MbTEC$^{FL}$ vs MbTEC$^{32K}$ and MbTEC$^{FL}$ vs MbTEC$^{PH}$ ) or 20 pmol (MbTEC$^{FL}$ vs MbTEC$^{FL}$ Y579D R581E) for each respective condition. For experiments with MbTEC$^{32K}$ and MbTEC$^{PH}$ reactions were initiated by the addition of 16.6 µL of $D_2O$ buffer (20 mM HEPES pH 7.5, 150 mM NaCl, 1mM TCEP, 95.1% D2O (V/V)) to 3 µL of protein and 0.4 µL of $MgCl_2$ + ADP (final D2O concentration of 78.9%, final $MgCl_2$ concentration of 2mM, final ADP concentration of 1 mM). Reactions proceeded for 3s at 0°C, and 30s, 300s and 3000s at 20°C before being quenched with ice cold acidic quench buffer, resulting in a final concentration of 0.6M guanidine HCl and 0.9% formic acid post quench. The exchange reactions for MbTEC$^{FL}$ vs MbTEC$^{FL}$ Y579D R581E were initiated by the addition of 13.5 µl of $D_2O$ buffer (20 mM HEPES pH 7.4, 150 mM NaCl, 1 mM TCEP 95.1% $D_2O$ (V/V)) to 2.5 µL of protein and 4.0 µL of $MgCl_2$ +ADP (final $D_2O$ concentration of 64.2%, final $MgCl_2$ concentration of 2 mM, final ADP concentration of 1 mM). Reactions proceeded for 3 s at 0°C, and 3 s, 30 s, 300 s and 3000 s at 20°C before being quenched with ice cold acidic quench buffer, resulting in a final concentration of 0.6 M guanidine HCl and 0.9% formic acid post-quench. All conditions and timepoints were created and run in independent triplicate. Samples were flash frozen immediately after quenching and stored at -80°C until injected onto the ultra-performance liquid chromatography (UPLC) system for proteolytic cleavage, peptide separation, and injection onto a QTOF for mass analysis, described below.

##### Protein Digestion and MS/MS Data Collection

Protein samples were rapidly thawed and injected onto an integrated fluidics system containing a HDx-3 PAL liquid handling robot and climate-controlled (2°C) chromatography system (LEAP Technologies), a Waters Acquity UPLC I-Class Series System, as well as an Impact HD QTOF Mass spectrometer (MbTec$^{FL}$v MbTec$^{32K}$or MbTec$^{FL}$ v MbTec$^{PH}$) (Bruker) or an Impact II QTOF Mass spectrometer (MbTec$^{FL}$ v MbTec$^{FL}$ Y579D R581E). The full details of the automated LC system were previously described in (13). The samples were run over an immobilized pepsin column (Affipro: AP-PC-001) at 200 µL/min for 4 min at 2°C. The resulting peptides were collected and desalted on a C18 trap column (Acquity UPLC BEH C18 1.7 µm column (2.1 × 5 mm); Waters 186004629). The trap was subsequently eluted in line with an ACQUITY 300Å, 1.7 µm particle, 100 × 2.1 mm BEH C18 UPLC column (Waters), using a gradient of 3-10% B (Buffer A 0.1% formic acid; Buffer B 100% acetonitrile) over 1.5 minutes, followed by a gradient of 10-25% B over 4.5 minutes, followed by a gradient of 25-35% B over 5 minutes, finally after 1 minute at 35% B a gradient of 35-80% B over 1 minute was used. Mass spectrometry experiments acquired over a mass range from 150 to 2200 m/z using an electrospray ionization source operated at a temperature of 200°C and a spray voltage of 4.5 kV.

##### Peptide identification

Peptides were identified from the non-deuterated samples of full length MbTEC using data-dependent acquisition following tandem MS/MS experiments (0.5 s precursor scan from 150-2000 m/z; twelve 0.25 s fragment scans from 150-2000 m/z). MS/MS datasets were analysed using FragPipe v18.0 (MbTec$^{FL}$v MbTec$^{32K}$or MbTec$^{FL}$ v MbTec$^{PH}$) or FragPipe v23.1 (MbTec$^{FL}$ v MbTec$^{FL}$ Y579D R581E) peptide identification was carried out by using a false discovery-based approach using a database of purified proteins and known contaminants (7, 9, 14). MSFragger was utilized, and the precursor mass tolerance error was set to -20 to 20ppm. The fragment mass tolerance was set at 20ppm. Protein digestion was set as nonspecific, searching between lengths of 4 and 50 aa, with a mass range of 400 to 5000 Da.

##### Mass Analysis of Peptide Centroids and Measurement of Deuterium Incorporation

HD-Examiner Software (Sierra Analytics) was used to automatically calculate the level of deuterium incorporation into each peptide. All peptides were manually inspected for correct charge state, correct retention time, appropriate selection of isotopic distribution, etc. Deuteration levels were calculated using the centroid of the experimental isotope clusters. Results are presented as relative levels of deuterium incorporation and the only control for back exchange was the level of deuterium present in the buffer (78.9% or 64.2%). Differences in exchange in a peptide were considered significant if they met all three of the following criteria: ≥5% change in exchange, ≥0.4 Da difference in exchange, and a p value <0.01 using a two tailed student t-test. Samples were only compared within a single experiment and were never compared to experiments completed at a different time with a different final $D_2O$ level. The data analysis statistics for all HDX-MS experiments are in are in the Source Data File according to published guidelines (15). The mass spectrometry proteomics data have been deposited to the ProteomeXchange Consortium via the PRIDE partner repository (12) with the dataset identifier PXD069610.

#### Fluorescence microscopy

##### Sample preparation

NIH/3T3 fibroblasts were cultured according to the manufacturer's instructions (ATCC). Briefly, cells were cultured in DMEM (Gibco) supplemented with 10 % FBS (Sigma-Aldrich) and Pen/Strep (Fisher Scientific) and passaged every 2 days. For transient transfection, they were seeded sub-confluently and co-transfected with two plasmids encoding mCherry-PH domain or eGFP-FL WT on the following morning with Lipofectamine 300 according to manufacturer's instructions. After 7 h of recovery, the media was replaced to DMEM without FBS for overnight starvation until microscopy imaging. During imaging in FluoroBrite media (Gibco), the media was spiked with 100 ng/mL of PDGFαβ (Gibco).

##### Data acquisition

Cells were imaged using a Zeiss LSM-980 confocal microscope with the Airyscan 2 module. Acquisition settings were chosen to satisfy the necessary 2x Nyquist sampling requirement for Airyscan imaging. Three-dimensional stacks were acquired over 40 time points with 30-second intervals. The integrated Airyscan 2 multiplexing functionality was employed to achieve faster imaging speeds. Maximum intensity projections of the Z-stacks were subsequently generated for visualization.

##### Image analysis

Representative regions of interest, where strong responses were observed, were first extracted from raw files. Analysis was then performed within these ROIs. Raw images were first denoised using a 5x5 median filter, followed by background subtraction via white-tophat transform with a rolling ball-disk (radius 7px). As MbTEC fluorescence was observed to accumulate in tube-like structures, a Frangi vesselness filter (**α**: 1.0, **γ**: 5.0, **σ**: 6, 8, 10, 12) was applied to both channels to pronounce any filamentous structures.

The frangi filter is a multi-scale, Hessian based ridge-detection algorithm, highlighting tubular structures, while suppressing "blobs" or plate-like features. Three frangi-filtering parameters were specified for this analysis:

- **α:** ("plate sensitivity") specifies how strongly plate-like structures are weighed down
- **γ:** ("structuredness") governs overall noise tolerance, dampening out low contrast regions
- **σ**: specifies the scale at which the gaussian filter during vessel detection is applied. Selecting a range of sigmas leads to the filter running across multiple scales, while the strongest response across all sigma values is captured.

The two frangi-filtered images were then averaged – resulting in a combined tubeness-response image for both channels. Segmentation was then performed by applying a global Otsu threshold scaled by a factor 0.5, and small objects were filtered ($<200\ px^2$). From these binary masks, features were extracted from the unprocessed image in both channels (area, mean intensity, median intensity). To account for photobleaching during image acquisition, the overall decrease in fluorescence was extracted from the raw files. Extracted intensities from the masked regions were then fitted to the obtained bleaching-curve and normalized to the first timepoint.

#### Preparation of $PIP_3$-containing nanodiscs

##### MSP1D1 expression and purification

MSP1D1 was purified as previously reported with additional anion exchange chromatography and gel filtration steps. Briefly, the MSP1D1 sequence was derived from a plasmid obtained from AddGene (ID: 173482) and cloned into pET28a vector with a N-terminal His-tag. The plasmid was transformed into *E.coli* BL21 Star. The cells were grown in LB supplemented with 100 ug/mL ampicillin at 37 °C and 180 rpm to an $OD_{600}$ of 0.7. Expression was induced with 0.2 mM IPTG and the culture was incubated overnight at 18 °C and 180 rpm.

The cells were harvested by centrifugation at 4000 rpm for 30 min and frozen in liquid nitrogen. The pellet was resuspended in lysis buffer (50 mM Tris pH 8.0, 400 mM NaCl, 5% glycerol, 1 mM $MgCl_2$, 0.25 % CHAPS, 5 mM imidazole, 2 mM DTT) supplemented with 250 U of Denarase (c-LEcta) and sonicated three times at 50% for 3 min with 1 sec on and off pulse. The lysate was cleared by centrifugation at 18 000 rpm for 30 min and loaded onto a $Ni^{2+}$-NTA column (HisTrap 5 mL FF, Cytiva) pre-equilibrated in 50 mM Tris pH 8, 400 mM NaCl, 5% glycerol, 5 mM imidazole, 2 mM DTT. The column was washed with 10 column volumes of 50 mM Tris pH 8, 400 mM NaCl, 5% glycerol, 20 mM imidazole, 2 mM DTT and followed by a step-gradient to 50 mM Tris pH 8, 400 mM NaCl, 5% glycerol, 500 mM imidazole, 2 mM DTT for elution of MSP1D1. The sample was dialyzed overnight against 50 mM Tris pH 8.0, 100 mM NaCl, 5% glycerol, 1 mM DTT, while simultaneously cleaving-off the His-tag using His-TEV protease (in-house). The dialyzed sample was loaded over a $Ni^{2+}$-NTA column (HisTrap 5 mL FF, Cytiva) for removal of the cleaved tag and protease. The flow-through was subjected to anion exchange chromatography (HiTrap Q 1 mL HP, Cytiva), and subsequent gel filtration using a 16/600 Superdex 75 pg column pre-equilibrated in 50 mM Tris pH 8, 100 mM NaCl, 5% glycerol, 1 mM TCEP. The protein was concentrated, snap-frozen in liquid nitrogen, and stored at -70 °C until further use.

##### Preparation of nanodiscs containing $PIP_3$

Nanodiscs were prepared as previously reported (16) with lipids purchased from Avanti Polar Lipids. Briefly, chloroform stocks of cholesterol (5 mg/mL), DOPC (5 mg/mL), DOPS (10 mg/mL), and DOPE (5 mg/mL) were prepared, whereas $PIP_3$ was solubilized in a chloroform:methanol:water (1:2:0.8) mixture. Lipids were combined in a borosilicate glass tube (Duran) at the following molar ratio: 20% cholesterol, 15 % DOPS, 20% DOPC, 35% DOPE and 10% $PIP_3$. The lipids were dried under a nitrogen stream, incubated under vacuum for 2 hours and rehydrated in buffer A (50 mM Tris pH 8, 100 mM NaCl, 1 mM TCEP). The mixture was freeze-thawed 5x with vortexing and sonication steps in between (Bandelin Sonorex Digiplus, 90% for 5 min at RT). The sample was supplemented with 0.5% dodecyl maltoside (DDM) in buffer A and incubated at room temperature for 1 hour. Next, Buffer A and MSP1D1 were added to obtain a MSP1D1:lipid molar ratio of 1:60 with a final concentration of 0.1% DDM. The mixture was incubated on ice for 1 hour, followed by gentle rotation overnight at 4°C on BioBeads SM-2 Absorbents (BioRad #1523920) for removal of the detergent. The nanodiscs were purified on a Superose 6 Increase 3.2/100 column and stored at 4°C until further use.
